# Mimicking two posttranslational modifications associated with oxidative stress affords phase separation of vimentin

**DOI:** 10.64898/2026.08.06.743216

**Authors:** Diego Moneo-Corcuera, Paula Martínez-Cenalmor, Alma E. Martínez, Dolores Pérez-Sala

## Abstract

Biomolecular condensates are membraneless compartments critical for the functional organization of cellular macromolecules in essential processes such as cell division, gene transcription or stress responses. We previously reported that vimentin filaments remodel into phase separated biomolecular condensates upon oxidative stress. This process requires vimentin single cysteine, C328, suggesting the involvement of oxidative modifications of this residue. Here, we aimed to generate vimentin condensates by inserting mutations mimicking posttranslational modifications associated with oxidative stress. In vimentin deficient cells, a cysteine oxidation mimetic mutant, vimentin C328D, formed only elongated particles or short filaments that evolved towards droplets upon serum deprivation or treatment with the oxidant diamide. Among vimentin posttranslational modifications rapidly responding to these stimuli, glycosylation confers filament stability whereas phosphorylation promotes disassembly. We observed that the O-deglycosylation inhibitor thiamet G, and the kinase inhibitors staurosporine and H-89, attenuated diamide-elicited vimentin C328D droplet formation, suggesting a potential glycosylation/phosphorylation interplay in this effect. Indeed, introducing phosphomimetic residues at certain single vimentin glycosylation and/or phosphorylation sites induced the formation of droplets, only if combined with the C328D mutation. In particular, the vimentin S49D,C328D mutant formed condensates that were reversibly dispersed by dilution through hypotonic shock. Therefore, mimicking C328 oxidation and S49 phosphorylation was sufficient to elicit vimentin phase separation. In vitro, purified vimentin S49D,C328D polymerized into a mixture of aberrant filaments and aggregates, which, in the presence of crowders, evolved towards paracrystals or clusters of beaded assemblies depending on pH. These findings highlight the role of C328 perturbations in the formation of biomolecular condensates and suggest a modulatory role of glycosylation/phosphorylation, thus shedding light on the processes regulating vimentin phase separation.

## Introduction

Tight regulation of the organization of cellular components is essential for cell function. Biomolecular condensates are membraneless compartments in which certain biomolecules (proteins, nucleic acids, etc.) concentrate through a phase separation phenomenon [1]. Biomolecular condensates thus constitute specialized environments where biological processes can be favored by the proximity of their constituents. Examples of cellular structures involved in key physiological functions that are formed through biomolecular condensation include the nucleolus, centrosomes and the nuclear pores [2–5]. In turn, aberrant phase transition can lead to disease, as evidenced in cancer, infectious diseases and neurodegenerative disorders [6, 7].

In proteins, the presence of multivalent surfaces together with intrinsically disordered domains capable of establishing multiple weak interactions is critical for phase separation [3]. This process is modulated by multiple factors, including concentration of the components, environmental conditions, such as pH or the presence of metals, nucleic acids or small molecules, or even cellular surfaces [8, 9]. Posttranslational modifications (PTMs) are decisive factors in phase separation due to their ability to change protein charge, hydrophobicity, conformation, and therefore, molecular interactions [10]. Nevertheless, the precise consequences of PTMs in protein condensation depend on the protein species. Thus, phosphorylation can favor or preclude phase separation depending on the substrate, whereas modifications of cysteine residues can modulate condensate formation in a structure dependent manner [11, 12].

Intermediate filaments are essential components of the cell cytoskeleton. They constitute a broad family of proteins that are the products of more than 60 genes, the expression of which is tightly regulated in a tissue specific manner [13]. Beyond providing structural support and mechanical resilience, intermediate filaments contribute to organelle homeostasis and integration of cellular functions, such as cytoskeletal crosstalk and response to stress [14]. These multiple roles are sustained by their sophisticated modulation through PTMs, which contribute to generate proteoforms that can associate in different assemblies [14, 15]. Intermediate filament monomers share a modular structure with disordered N-terminal (head) and C-terminal (tail) domains, flanking a central α-helical rod domain [16]. Interestingly, the isolated head domains of certain intermediate filaments can undergo phase separation [17], whereas in the context of the filament they can engage in multiple labile interactions that contribute to the modulation of filament assembly [18, 19]. In fact, under certain conditions, filaments can dissociate yielding droplets [12, 19].

Vimentin is a type III intermediate filament protein mainly expressed in the cytoplasm of mesenchymal cells, where it plays key roles in mechanotransduction, resistance to various types of stress, cell signaling and cytoskeletal crosstalk, with consequences in cell migration, division and pathogen invasion [20, 21]. Interestingly, vimentin can form different types of assemblies, including filaments, bundles, squiggles, droplets and several extracellular forms, in response to cellular or subcellular conditions [21–23]. The ability of vimentin to remodel into widely different assemblies relies on a delicate balance between PTMs and context factors [14, 24]. Vimentin constitutes a hub for PTMs, and more that 150 PTM sites have been identified along its sequence. Moreover, at least 37 phosphorylation sites have been reported within the first 117 residues of human vimentin, according to numerous cellular and proteomic studies [25–27] and to the information available in the PhosphositePlus database (https://www.phosphosite.org/proteinAction.action?id=2622&showAllSites=true) [28]. Importantly, vimentin and other type III intermediate filament proteins, such as GFAP and desmin, possess a single cysteine residue, C328 in vimentin, that is redox sensitive, and is a hot spot for PTMs [29, 30]. More than 15 different modifications have been reported for vimentin C328 (reviewed in [14]). Certain modifications have been identified under basal or control conditions, and a switch to other modifications can occur upon exposure to various stresses or environmental factors [27]. Remarkably, C328 modification by structurally diverse moieties has distinct functional consequences on protein assembly [31]. The complexity and precision of this regulation can be envisaged if we consider that the model of the vimentin filament contemplates 40 monomers per cross-section (ten tetramers) [32], with cysteine residues from two adjacent tetramers appearing in relative proximity [30, 32]. Moreover, the PTMs present in every monomer may be different, and could occur in different combinations, giving rise to a plethora of proteoforms. In addition, modifications can also vary depending on the phase of the cell cycle or the subcellular localization of the filaments, in order to control the spatiotemporal functions of the network in processes such as cell migration or division [26, 33–35].

In earlier works, we reported the ability of cellular vimentin filaments to extensively remodel into droplets in response to oxidants, in a manner dependent on the presence of C328 [36, 37]. Recently, we demonstrated that oxidant elicited vimentin droplets behave as biomolecular condensates [12], and showed that C328 acts as a checkpoint for phase separation under these conditions. Nevertheless, given the complexity of vimentin PTMs, it is not known whether C328 modification is sufficient for the transition of filaments into droplets in response to oxidants. The functions of vimentin condensates could be multiple and have been proposed to act as nucleating centers for filament elongation, promote the “wetting” and protection of actin fibers and preserve the protein under oxidative stress [12, 18, 38]. However, the transient, dynamic and likely complex nature of these structures imposes limitations for their study.

In this work, we have attempted to pinpoint residues and modifications that could be critical for vimentin droplet formation in response to oxidative stress, taking advantage of the possibility of generating PTM mimetics by site directed mutagenesis. Our results confirm the requirement for C328 in oxidant induced phase separation of vimentin, and suggest that C328 oxidation, together with a selective single phosphorylation at the vimentin head domain could be sufficient to drive this process, which may be further regulated by factors such as pH and biomolecule agglomeration.

## Materials and Methods

### Reagents

Diamide, 1,6-hexanediol, 1,5-hexanediol, methoxypolyethylene glycol maleimide (MALPEG-5, with PEG number average molecular weight of 5000), 4,6-diamidino-2-phenylindole (DAPI), H-89 and staurosporine were from Merck. Thiamet G was from Santa Cruz Biotechnology. Y27632 was from Calbiochem. Antibodies used were anti-vimentin clone V9 and its Alexa-488 conjugated form (Santa Cruz Biotechnology, sc-6260), anti-vimentin D21H3 (Cell Signaling), and anti-actin polyclonal antibody from Sigma. Oligonucleotides for site directed mutagenesis were obtained from IDT. Dextran 500, diamide and dibromobimane (DBB) were from Sigma and bis-maleimide hexane (BMH) from ThermoFisher.

### Cell culture and treatments

SW13/cl.2 adrenal carcinoma cells, deficient in cytoplasmic intermediate filaments, were the generous gift of Prof. A. Sarriá (University of Zaragoza, Spain). A549 cells depleted of vimentin by CRISPR-Cas9 have been previously described [35]. MEF wt and *Vim*^−/−^ were the gift of Prof. J. Eriksson (Abo Academy, Finland). SW13/cl.2 cells and MEF were cultured in DMEM high glucose, supplemented with 10% (v/v) fetal bovine serum and antibiotics (100 U/ml penicillin, 100 μg/ml streptomycin), all from Gibco. SW13/cl.2 cells stably expressing vimentin wt were previously reported [36]. Stably transfected cells were cultured in the presence of the selection antibiotic, G-418 (Gibco). Unless otherwise stated, treatments were performed in serum free medium. Treatment with diamide was generally carried out by incubation at 37 °C with 1 mM for 15 min. Cysteine crosslinkers DBB and BMH were added to cells at 100 µM for 15 min before the addition of diamide. Incubation with thiamet G at 10 µM was carried out for 15 min and it was added at the same time than diamide. H-89 and staurosporine were added at 10 µM and 50 nM, respectively, 5 min before diamide addition. To assess the effect of aliphatic alcohols, cells were treated in the absence or presence of 1 mM diamide for 20 min, and alcohols were added at 3.3% (w/v) final concentration for the last 5 min of the incubation. Hypotonic shock was induced by incubating cells with culture medium diluted 1:10 in water. Hypertonic shock was achieved by incubation with culture medium supplemented with 150 mM NaCl.

### Plasmids and transfections

The bicistronic plasmids pIRES2 DsRed-Express2 vimentin (RFP//vimentin) for expression of the red fluorescent protein DsRed-Express2 and human vimentin wt as separate products, and the GFP-vimentin fusion construct have been reported previously [36]. Mutants employed in this study were generated by site directed mutagenesis employing the NZyTech mutagenesis kit, following the instructions of the manufacturer, and oligonucleotides specified in Suppl. Table 1. Oligonucleotides for vimentin C328D and C328W mutants have been previously reported [31]. For cell transfection, Lipofectamine 2000 (Invitrogen) was routinely used. In a typical transfection, cells in a p35 dish were incubated with a mixture containing 1 µg of DNA and 3 µl of Lipofectamine 2000, prepared in Optimem (Invitrogen), for 5 h in culture medium without antibiotics. Subsequently, the transfection mixture was removed, and cells were allowed to recover for 48 h in medium without antibiotics, before being used for experiments.

### SDS-PAGE and western blot

After treatment under the various experimental conditions, cell monolayers were washed with cold PBS. Cells were lysed by gentle scraping and forced passes through a narrow needle (26 1/2G) in lysis buffer containing: 50 mM Tris-HCl, pH 7.5, 0.1. mM EDTA, 0.1 mM EGTA, 0.5% (w/v) SDS, 0.1 mM ꞵ-mercaptoethanol, plus 50 mM NaF, 20 mM sodium orthovanadate and protease inhibitors aprotinin, leupeptin, trypsin inhibitor and pepstatin, each at 2 µg/ml, and Pefablock at 1.3 mM. Lysates were either precleared by centrifugation at 10000 g for 5 min at 6 °C, or directly mixed with Laemmli sample buffer and incubated at 95 °C for 5 min. Protein concentration in lysates was determined with the BCA kit from Pierce. Routinely, aliquots containing 20 µg of protein were separated on 10% (w/v) polyacrylamide gels. Proteins were transferred onto Immobilon P membranes (Millipore) using a semi-dry transfer unit from Bio-Rad, and a three-buffer system, following the instructions of the manufacturer. Blots were blocked by incubation with 2% (w/v) non-fat evaporated milk and incubated with the antibodies of interest at 1:1000 dilution for primary and 1:2000 dilution for secondary HRP-conjugated antibodies, at room temperature for 1 h. Signals were obtained using the ECL chemiluminescence system and exposure of blots to Hyperfilm (Cytiva), which were developed in an Agfa Curix 60 processor.

### Assessment of C328 occupancy

To estimate the extent of C328 modification after the various treatments, several different strategies were used. In a set of experiments, cells were treated in the absence or presence of diamide for 15 min, after which, they were incubated with two different cysteine bifunctional crosslinkers, namely DBB and BMH, at 100 µM final concentration for additional 15 min. At the end of the treatment, cells were lysed and lysates were analyzed by gel electrophoresis under non reducing and reducing conditions. Vimentin was detected by western blot. In this assay, modification of vimentin cysteine upon treatment with diamide results in lower availability of the thiol group for subsequent crosslinking by the cysteine reagents. In a second set of assays, the presence of vimentin with available thiol group was detected by its complexation with MALPEG-5, which results in an upward shift in the electrophoretic mobility of the protein band. Lysates from control and diamide treated cells were obtained as above and aliquots containing 1 mg/ml of total protein were incubated with 1 mM MALPEG-5 for 15 min at room temperature before being analyzed by gel electrophoresis under non reducing or reducing conditions, followed by western blot for the detection of vimentin. As a third approach, cells were treated in the absence or presence of diamide and lysed in cold lysis buffer without reducing agents. Immediately, lysates were incubated with magnetic beads coupled to maleimide. In this assay, vimentin with free thiol group is retained in the beads, whereas vimentin with modified C328 appears in the unbound fraction. The levels of vimentin in the input and the unbound fraction were estimated by western blot, from which the proportion of vimentin with non-available (modified) thiol group can be calculated. No vimentin was detected in the eluate after washing the beads with excess DTT, thus ruling out the potential retention of forms modified by persulfidation or by mixed disulfide with species containing additional thiol groups.

### Immunofluorescence and confocal microscopy

Cells grown on coverslips (Epredia) or in glass bottom p35 dishes (Mattek Corporation) were washed with PBS and fixed by incubation for 25 min at room temperature with 4% (w/v) paraformaldehyde (PFA). After three washes with PBS, cells were permeabilized by incubation in 0.1% (v/v) Triton X-100 for 20 min at room temperature, and blocked by incubation with 1% (w/v) BSA in PBS. Antibodies were routinely used at 1:200 dilution in blocking solution, and incubations were carried out at room temperature. Nuclei were counterstained with DAPI at 3 µg/ml in blocking solution. Coverslips were mounted with Fluorsave (Calbiochem). Immunostained cells were visualized on Leica SP5 or SP8 confocal microscopes. Individual sections were taken every 0.5 µm, with an oil immersion objective at 63x magnification. Single confocal sections or overall projections are shown as indicated. For visualization of live cells, a thermostatized chamber was used. Scale bars are 20 µm, except when specified otherwise.

### FRAP assays

Cells seeded on glass bottom cell culture dishes (Mattek Corporation) were transfected with a mixture of RFP//vimentin S49D,C328D plus GFP-vimentin S49D,C328D plasmids, in a 4:1 proportion. Forty eight hours after transfection, cells were monitored in a Leica SP5 microscope equipped with a thermostatized support. Briefly, after acquiring a pre-bleach image with a zoom of 3x, a region of interest (ROI) of 10 x 2.8 µm was bleached by three pulses of the 488 nm laser line at 80% potency. A postbleach image was immediately obtained and then single sections were acquired every five seconds during three minutes. Fluorescence intensity of the ROI from these assays was quantitated with LasX software.

### Protein preparation

Purified recombinant human vimentin wild-type (wt) and vimentin S49D,C328D mutant, were obtained from Abvance Biotech S.L. (Madrid, Spain). EDTA removal and protein refolding were performed as previously described [39]. Briefly, purified proteins in 8 M urea, 5 mM Tris-HCl (pH 7.6), 1 mM EDTA, 10 mM β-mercaptoethanol, 0.4 mM PMSF and approximately 150–200 mM KCl were first diluted 1:10 with EDTA-free buffer and concentrated down to the initial volume by ultrafiltration using Amicon Ultra centrifugal filter units (10 kDa molecular weight cutoff, Millipore). This step was repeated twice. Protein refolding was performed by stepwise dialysis against 5 mM PIPES-Na (pH 7.0) containing 1 mM DTT and decreasing concentrations of urea (6, 4, and 2 M), followed by two additional dialysis steps in urea-free 5 mM PIPES-Na (pH 7.0) containing 0.25 mM DTT. The final dialysis was carried out for 16 h at 16 °C. Refolded protein preparations were clarified by ultracentrifugation at 120,000 × g for 15 min at 4 °C, and the supernatants were aliquoted and stored at −80 °C until use. Protein concentration was estimated from its absorbance at 280 nm, using an extinction coefficient of 22450 M^−1^ cm^−1^. Throughout this work, amino acid numbering for vimentin includes the initial methionine.

### In vitro polymerization of vimentin

For in vitro assays, soluble oligomeric species of vimentin wt or S49D,C328D mutant at 0.2 mg/ml in 5 mM PIPES-Na, 0.1 mM DTT, were subjected to polymerization by incubation in the presence of 150 mM NaCl for 1 hour at 37 °C. The final pH of the incubation mixture was 7.0 or 7.5, as indicated. When specified, macromolecular crowding conditions were established by adding dextran 500 at 150 mg/ml, final concentration. Stock solutions of dextran 500 were prepared by dialysis in 5 mM PIPES-Na, 0.1 mM DTT, at the corresponding experimental pH (7.0 or 7.5). The precise concentration of the crowder stock solutions was determined from the increment in the refractive index (0.145 ml/g for dextran 500).

### Electron microscopy

Carbon support grids (MESH CF 400 CU UL, Aname) were placed onto drops of 0.1% glutaraldehyde-fixed mixtures, subsequently washed with water, and negatively stained with 2% (w/v) uranyl acetate (Merck) for 60 s. In experiments in which vimentin S49D,C328D was polymerized in the presence of dextran 500 at pH 7.5, protein structures exhibited poor adherence to the carbon support grid. In these cases, samples were sedimented onto the grids by centrifugation at 12,000 rpm for 5 min before negative staining. Introducing a similar centrifugation step during processing of vimentin wt samples did not alter the structures observed. Grids were inspected on a JEOL JEM-1230 transmission electron microscope operating at 100 kV and equipped with a CMOS TVIPS TemCam-F416 digital camera. Images were acquired at 45 K magnification and the image analysis was performed using FIJI.

### Image analysis

For analysis of confocal microscopy images, either the FIJI or the LAS X software were used. The “analyze particles” command (FIJI) was used to obtain number of particles, their area, circularity and perimeter. The correlation coefficient (the ratio between the standard deviation of fluorescence intensity and the average fluorescence intensity obtained with FIJI) for individual cells was used as an inverse index of the degree of dispersion of vimentin droplets. Particle motility was assessed with the FIJI “tracking particles” plugin. For electron microscopy images, the width of filamentous structures, the diameter of particles, and the length of the axial periodicity of paracrystal-like assemblies were measured manually using the “straight line” tool after calibrating each image with the electron microscopy scale bar. Measurements were obtained from at least three independent experiments.

### Statistical analysis

All experiments were repeated at least three times. Data were analyzed with GraphPad v9 software. Data following a normal distribution were analyzed by parametric one-way ANOVA followed by Tukey test for multiple comparisons, or the Student t-test for comparison between two experimental conditions. For data not adjusting to a normal distribution, the non-parametric Kruskal-Wallis test, followed by Dunn’s test was employed for multiple comparisons, whereas pairs of datasets were compared with the Mann-Whitney test. Data are generally presented as average values ± SEM. When specified, data are represented as box plots, with the central mark depicting the median, the edges of the box corresponding to the 25^th^ and 75^th^ percentiles, and whiskers extending to the most extreme data points. Statistical significance was defined as *p<0.05, **p<0.01; ***p<0.001; ****p<0.0001.

## Results

### Effect of oxidant treatment on the availability of the vimentin C328 thiol

We have previously shown that treatment of cells expressing vimentin wt with various oxidants elicits the remodeling of vimentin filaments into droplets with the characteristics of biomolecular condensates [12]. This is illustrated for the oxidant diamide in Fig. 1A. This process requires the presence of C328, since filaments formed by vimentin C328A, C328S or C328H mutants are resistant to this effect [12, 31, 36]. These observations strongly suggest that the modification of C328, likely by oxidation, is involved in vimentin phase separation. Here, we have obtained further insight into this process. The landscape of C328 PTMs can be highly complex, since it can comprise small or bulky moieties, as well as modifications potentially altering the charge of this site (schematized in Fig. 1B). Bulky modifications of C328 include disulfide bond formation with other proteins, such as other vimentin monomers [24, 37], or incorporation of other molecules, as in certain forms of lipoxidation, glutathionylation and CoAlation [27, 40–43]. In turn, small oxidative modifications may include nitrosylation, sufenylation, sulfinylation and sulfonylation [27, 42, 44]. In addition, some of these modifications such as sulfinylation, glutathionylation or CoAlation introduce negative charges at the site of the modified cysteine. Moreover, at any given time, cysteine residues from different vimentin monomers can bear diverse PTMs in variable proportions [27].

**Figure 1.**
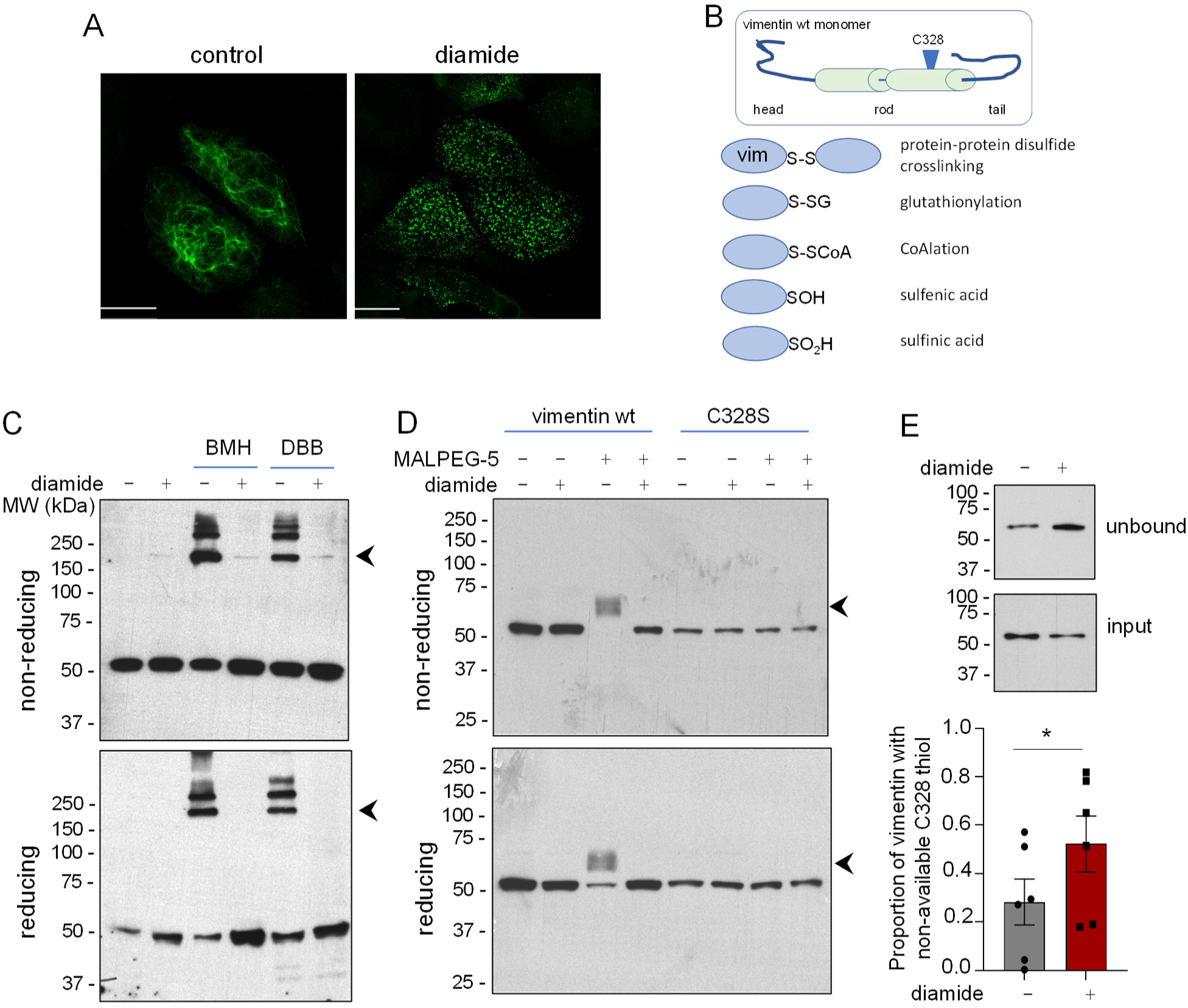
Modifications of vimentin C328 occurring upon oxidative stress elicited by diamide. (A) Remodeling of vimentin filaments into droplets by diamide treatment. SW13/cl.2 cells stably expressing vimentin wt were treated in the absence or presence of 1 mM diamide for 15 min, and the remodeling of vimentin was assessed by immunofluorescence and confocal microscopy as detailed in Methods. (B) Potential modifications of vimentin C328 occurring upon diamide treatment. A cartoon of the vimentin monomer shows the head, rod and tail domains, along with the approximate position of C328 (modified from [22]). In the lower panel, vimentin is represented by blue ovals on which potential modifications affecting the C328 thiol upon treatment of cells with diamide are schematized. (C) Diamide treatment blocks the formation of vimentin crosslinks by bifunctional cysteine crosslinkers and induces disulfide crosslinking on its own. SW13/cl.2 transfected with vimentin wt were treated in the absence or presence of 1 mM diamide for 15 min, before the addition of the cysteine crosslinkers BMH or DBB at 100 µM, final concentration for 15 min. Cells were lysed without reducing agents and aliquots from the cell lysates were analyzed by gel electrophoresis under non-reducing (upper panel) or reducing (lower panel) conditions, followed by western blot with anti-vimentin antibody V9. Arrows mark the position of the products of cysteine crosslinking (resistant under reducing conditions); arrowheads mark the position of disulfide-bonded vimentin (reversed under reducing conditions). (D) Detection of vimentin with free thiol group by MALPEG-5 binding. SW13/cl.2 cells expressing vimentin wt or C328S were treated in the absence or presence of diamide, as indicated, after which, lysates were prepared in the absence of reducing agents, and immediately incubated with MALPEG-5 at 1 mM, final concentration, for 15 min at room temperature. Finally, lysates were analyzed by gel electrophoresis under non reducing and reducing conditions, and vimentin was detected by western blot. Arrowheads mark the position of the MALPEG-5-modified vimentin band, which displays an upward shift indicating that the C328 thiol group was available for modification. (E) SW13/cl.2 cells expressing vimentin were treated with 1 mM diamide for 15 min. Lysates from control and diamide-treated cells were incubated with maleimide-agarose to retain proteins with free thiol groups. Vimentin with modified C328 appeared in the unbound fraction. The graph shows the estimation of the proportion of vimentin with modified C328 depicted as mean values ± SEM of six independent assays. *p<0.05 by paired Student t-test.

In view of this complexity, we decided to assess the overall modification of vimentin C328 upon diamide treatment employing several complementary approaches. Firstly, we explored the ability of diamide to elicit vimentin disulfide crosslinking and to preclude crosslinking by other agents. We previously showed that vimentin is susceptible of crosslinking by bifunctional cysteine reagents in a manner dependent on the presence of C328 [36, 45]. Here we observed that treatment of cells expressing vimentin with the cysteine crosslinkers DBB or BMH promoted vimentin crosslinking, demonstrated by the appearance of oligomeric species, detected by western blot with anti-vimentin antibody, which were resistant under reducing conditions (Fig. 1C). Preincubation of cells with diamide precluded vimentin cysteine crosslinking by both agents, indicating a loss of availability or accessibility of the C328 thiol. Moreover, a small amount of vimentin oligomer could be detected in samples from cells treated with diamide, likely representing disulfide crosslinked vimentin, since it was reversed under reducing conditions (Fig. 1C, arrowheads).

Secondly, we employed a strategy consisting in the incubation of cell lysates with pegylated maleimide (MALPEG), which results in an upward shift in the electrophoretic mobility of proteins with free thiol groups [46]. Here, we observed that a pegylated maleimide of average size 5 kDa (MALPEG-5) elicited a clear shift in vimentin present in lysates from control cells, which was blunted in lysates from diamide treated cells (Fig. 1 D, arrowhead), indicating a lower availability of the C328 thiol after diamide treatment. Curiously, when analyzed under reducing conditions, a small proportion of vimentin from control cells did not display the MALPEG-5 elicited shift (Fig. 1D, lower panel). This could indicate that this proportion of vimentin is reversibly modified, for instance by low molecular weight dithiol containing species, in which case the shift induced by MALPEG-5 would be reversed under reducing conditions, or that there is a partial reversion of the maleimide thioether bond. As a control for this assay, we confirmed that MALPEG-5 did not alter the electrophoretic mobility of a vimentin C328S mutant.

Finally, we assessed the extent of C328 modification upon diamide treatment by subjecting lysates from control and diamide treated cells to denaturation and retention of proteins with free thiol groups on maleimide-conjugated beads. The presence of vimentin in the unbound fraction provides an index of C328 modification, since only the protein bearing a free thiol group is expected to be retained in the beads [24]. This assay yielded an estimate of 28 ± 8% and 69 ± 9.7% (mean ± SEM) of vimentin C328 modified in control and diamide-treated cells, respectively (Fig. 1E).

Altogether, these results indicate that diamide treatment significantly decreases vimentin C328 free thiol content at the expense of several modifications, which may include disulfide bond formation and several types of oxidation.

### Effect of perturbations at the 328 site of vimentin on filament formation

Next, we explored the potential impact of structural changes at C328 on filament assembly by introducing mutations at this site that either increase the size of the residue, namely, a cysteine to tryptophan mutation, or alter its charge by means of a cysteine to aspartic acid mutation (Fig. 2A). In fact, mutation to aspartate has been used as a mimetic of cysteine oxidation, for instance to sulfinic acid [31, 47], since, similar to the sulfinic group, the aspartate carboxylic group introduces a negative charge, although it does not possess the exact geometry [48]. Then, we assessed the assembly of these mutants in comparison to vimentin wt under normal growth conditions as well as in response to serum deprivation and diamide treatment (Fig. 2B). Vimentin wt was not detectably altered by serum deprivation and underwent the typical remodeling into droplets upon diamide treatment, as shown above (Fig. 2B, upper row images). The vimentin C328W mutant was able to form an extended filament network, with some filament bundles. Notably, this mutant was fully resistant to serum withdrawal and to the effect of diamide, as illustrated by the preservation of filaments in the presence of the oxidant (Fig. 2B, middle row images). Thus, the C328W mutation protected vimentin from diamide-eliciting remodeling, consistent with the thiol selectivity of diamide modification. Remarkably, the pseudo-oxidation mimic vimentin C328D mutant was unable to form long filaments, and only yielded small accumulations apparently interconnected by thin fibers, indicating that a negative charge at the site of C328 is not well tolerated for filament assembly and/or elongation (Fig. 2B, lower row). Serum deprivation partially decrased the interconnectivity of vimentin C328D structures, as indicated by an increase in the number of particles. Of note, vimentin C328D was still susceptible to diamide treatment, which increased particle circularity. These morphological changes are quantified in Fig. 2C and D. In addition, analysis of the three constructs by western blot with antibodies with epitopes located at the vimentin N-terminal (D21H3) or C-terminal (V9) domains, revealed that vimentin C328W and C328D yielded a main band compatible with the full length protein, which did not show apparent changes in electrophoretic mobility or integrity with respect to the wt (Fig. 2E). This suggests that the organization of vimentin C328D in small accumulations is due to an effect on assembly and not on the integrity of the protein.

**Figure 2.**
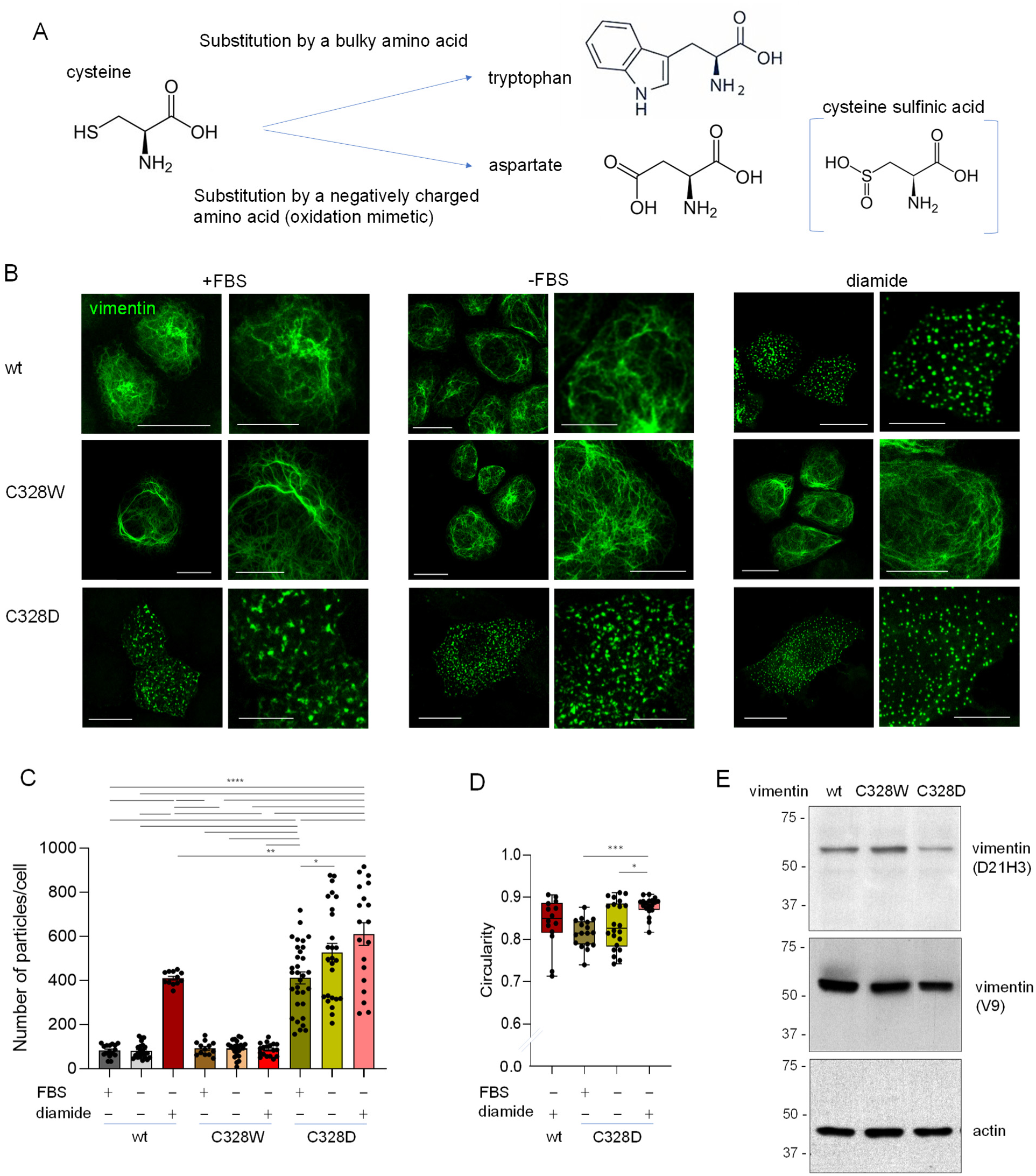
Impact of mutations at the 328 site of vimentin on droplet formation. (A) Scheme showing the structure of the cysteine residue and of the residues introduced at the 328 site to increase the size of the lateral chain (tryptophan) or to incorporate a negative charge (aspartate). In addition, the structure of a cysteine sulfinic acid is shown to highlight the similarity with aspartate. (B) Effect of introducing tryptophan or aspartate at the 328 site on vimentin assembly. SW13/cl.2 cells were transfected with the indicated constructs and the assemblies formed were monitored by immunofluorescence with anti-vimentin antibody under normal growth conditions (+FBS), upon serum deprivation for 15 min (-FBS) or after treatment with 1 mM diamide for 15 min in serum-free medium (diamide). The number of particles per cell (C) and their circularity (D) in the cells showing no filaments, was quantitated from at least three independent experiments totaling at least 13 cells per experimental condition for (C) and 14 for (D). Results are shown as mean ± SEM (C) or as box plots (D). Statistical significance was evaluated by one-way ANOVA followed by Tukey’s test (C) and Kruskal-Wallis followed by Dunn’s multiple comparisons test for (D). *p<0.05; **p<0.01; ***p<0.001; ****p<0.0001. (E) Lysates from cells transfected with the indicated constructs were analyzed by SDS-PAGE followed by western blot with antibodies against vimentin N-terminus (D21H3) or C-terminus (V9). Actin was used as a control.

### Potential role of vimentin deglycosylation and phosphorylation in diamide-elicited vimentin C328D droplet formation

The observation that serum deprivation and diamide treatment increased the number and circularity of vimentin C328D assemblies, respectively, suggests that the cysteine to aspartate mutation, mimicking an oxidized state of C328, does not fully recapitulate the features of diamide elicited droplets. Therefore, more disruptive modifications of C328 and/or additional modifications or interactions at the level of vimentin or other proteins, would be required to drive this process. Among modifications rapidly responding to serum availability and/or redox regulation, glycosylation and phosphorylation play an important role in the modulation of vimentin wt organization by stabilizing or destabilizing filaments, respectively (schematized in Fig. 3A) [25, 49, 50]. Therefore, we explored the effect of pharmacological modulators of these processes on vimentin C328D assemblies and their further remodeling into droplets by diamide. Thiamet G blocks O-deglycosylation by inhibiting the enzyme O-linked ꞵ-N-acetylglucosamine hydrolase (O-GlcNAcase or OGA) [51]. Treatment with thiamet G increased the connectivity and length of vimentin C328D structures, and prevented their remodeling by diamide, favoring the persistence of an irregular mesh of poorly defined filamentous structures (Fig. 3B). This suggests that promoting the presence of vimentin C328D in a glycosylated form increases its capacity to form interconnected structures, even in the presence of the oxidant. On the other hand, the relatively selective PKA inhibitor H-89 and the broad spectrum kinase inhibitor staurosporine, also favored the presence of vimentin C328D in small accumulations and short, linked filaments, and attenuated droplet formation by diamide (Fig. 3B). Nevertheless, the RhoK inhibitor Y27632 did not significantly mitigate the effect of diamide (Fig. 3B). These effects were quantitated by measuring the number (Fig. 3C), circularity (Fig. 3D) and perimeter (Fig. 3E) of vimentin C328D particles under the various experimental conditions. This showed that thiamet G, H-89 and staurosporine, but not Y27632, tended to decrease particle number in the absence and presence of diamide (Fig. 3C), and blocked the increase in circularity induced by the oxidant (Fig. 3D). Moreover, the same agents increased the perimeter of vimentin C328D assemblies in the presence of diamide (Fig. 3E). Taken together, these observations, summarized in Fig. 3F, suggest that O-deglycosylation and phosphoryIation may play a role in the formation of droplets elicited by diamide, and indicate that there may be some selectivity with respect to the kinases involved in these effects.

**Figure 3.**
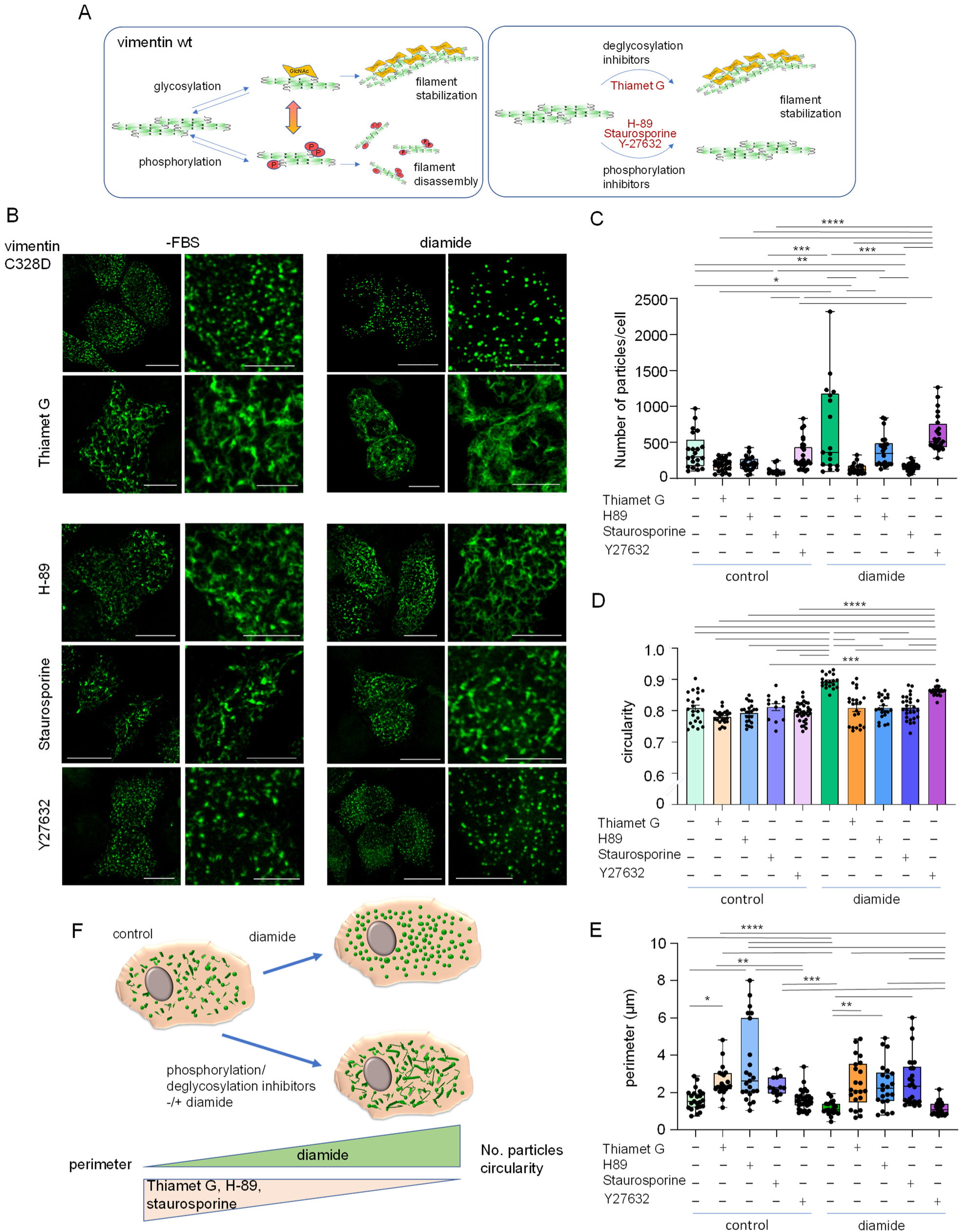
Effect of inhibitors of deglycosylation or phosphorylation on vimentin C328D assemblies. (A) Cartoon depicting the consequences of vimentin glycosylation and phosphorylation on filament stability (left panel), and the pharmacological tools employed to modulate these processes (right panel). Vimentin phosphorylation is known to contribute to filament disassembly, whereas glycosylation improves filament stability. These processes can be modulated by kinase inhibitors or deglycosylation inhibitors, both of which are expected to promote filament stabilization. (B) Impact of inhibitors of deglycosylation or phosphorylation on the assembly of vimentin C328D. SW13/cl.2 cells were transfected with the vimentin C328D construct. Forty eight hours after transfection, they were treated with 10 µM H-89 or 50 nM staurosporine for 5 min before or with 10 µM Y27632 for 15 min before the addition of 1 mM diamide for additional 15 min. When specified, thiamet G (10 µM) was added at the same time than diamide. Cells were fixed and vimentin C328D assemblies were monitored by immunofluorescence with anti-vimentin V9 antibody. (C-E) The number of particles (C), their circularity (D) and perimeter (E) were quantitated from a minimum of three experiments totaling at least 13 cells per experimental condition. The statistical analyses applied were the Kruskal-Wallis followed by Dunn’s multiple comparisons test (graphs C and E), and one-way ANOVA followed by Tukey’s test (graph D); ***p<0.001, ****p<0.0001. (F) Graphic summary of the morphological outcome of these assays.

### Specific phosphomimetic mutations favor droplet formation by vimentin C328D

Although O-deglycosylation and phosphorylation can occur throughout the cell and affect a myriad of proteins, we initially considered the possibility that the protective effects of inhibitors of these processes on droplet formation could be due to their impact on vimentin itself. The vimentin sequence presents numerous phosphorylation sites, most of which are located in the N-terminal domain, as depicted in Fig. 4A (left and center, modified from the PhosphositePlus database, https://www.phosphosite.org/proteinAction.action?id=2622&showAllSites=true) [28]. Among them, we and others have previously identified vimentin S49, S56 and S72 (Fig. 4A, center) as sites phosphorylated in conditions associated with oxidative stress [27, 52]. Notably, vimentin S49 is also an important target for glycosylation, involved in vimentin filament stability [49] (Fig. 4A, right). Therefore, an interplay or competition between glycosylation and phosphorylation could occur at this site. As a first approach to explore the potential contribution of vimentin phosphorylation to the effect of diamide on vimentin C328D, we generated double mutants introducing phosphomimetic residues at sites S49, S56 or S72, in combination with the C328D mutation, and assessed their behavior in cells (Fig. 4B-F). Of the three double mutants, the phospho/oxidation mimetic vimentin S49D,C328D was the construct that yielded droplets with the highest circularity under basal conditions in SW13/cl.2 cells, and the weakest response to diamide (Fig. 4B). In turn, vimentin S56D,C328D and S72D,C328D formed squiggle-like structures and slightly irregular dots, respectively, which in both cases evolved towards droplets upon diamide treatment (Fig. 4B). These results are quantitated in Fig. 4D. The different morphology of the assemblies formed by the three double mutants could also be observed in A549 *Vim*^−/−^cells, as well as in MEF *Vim*^−/−^ transfected with these constructs (Fig. 4C). In these cell types, S49D,C328D was also the construct that formed most circular structures (quantitated in Fig. 4E, F), which indicates that its tendency to form droplets is not cell type dependent. These results suggest that the functional consequences of phosphorylation of different sites in the vimentin head domain are not superimposable. Moreover, our observations indicate that, although oxidants can elicit a plethora of PTMs in vimentin, introducing PTM mimetic mutations at just two sites, particularly at S49 and C328, yields droplets resembling those elicited by diamide treatment of vimentin wt. However, the contributions of these two sites are different. Substitution of S49 by alanine or aspartate in vimentin wt did not preclude the formation of a filament network of normal appearance (Suppl. Fig. 1A). Therefore, perturbations at S49 are not sufficient to elicit vimentin remodeling. Moreover, both S49 mutants were susceptible to the effect of diamide, with the formation of droplets of high circularity, although they appeared slightly rounder in the case of the S49D mutant (quantitated in Suppl. Fig. 1B). Thus, it could be interpreted that these mutations do not hamper the impact of modifications at C328. However, in the context of the C328D mutation, the S49A substitution did not seem to enhance circularity, since an S49A,C328D vimentin mutant formed assemblies more interconnected and of lower circularity than those of vimentin S49D,C328D, and therefore more similar to those of vimentin C328D, both in the absence and presence of diamide (Suppl. Fig. 1C, quantitated in Suppl. Fig. 1D).

**Figure 4.**
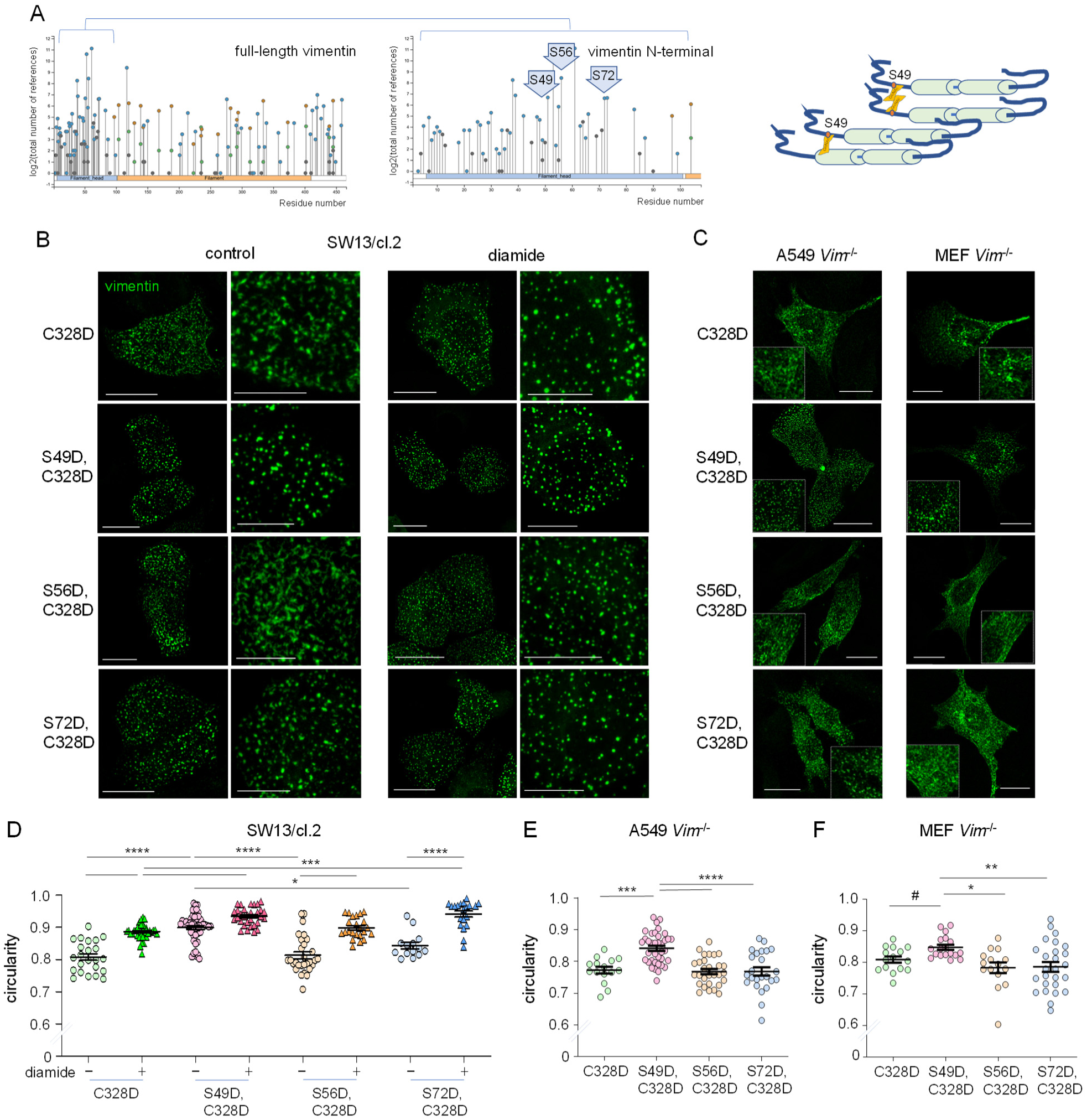
Importance of phosphorylation sites in droplet formation by vimentin C328D. (A) Schemes of phosphorylation sites reported for human vimentin. Plots have been obtained from the PhosphositePlus database (see text for details). The left image shows a lollipop plot of the PTM sites (blue circles, phosphorylation; green, acetylation; brown, ubiquitination; grey, other) reported along the whole vimentin sequence, whereas the center image shows a close up of the sites in the head domain. The residues of interest for our study are highlighted in block arrows. The cartoon at the right depicts the position of S49 and its potential involvement in glycosylation dependent stabilization of vimentin assembly. (B) Effect of introducing phosphomimetic mutations on vimentin C328D organization and response to diamide. SW13/cl.2 cells were transfected with the indicated constructs and 48 h later they were treated in the absence or presence of 1 mM diamide for 15 min. The organization of the mutant vimentin constructs was assessed by immunofluorescence. Regions of interest for every condition are enlarged in the images at right. (C) The organization of vimentin constructs was explored in A549 *Vim*^−/−^ cells (left images) or MEF *Vim*^−/−^ (right images) incubated for 15 min in serum free medium, by immunofluorescence with the V9 anti-vimentin antibody. Scale bars, 20 µm in main images and 10 µm in zoomed-in images. (D-F) The circularity of vimentin assemblies under the indicated experimental conditions has been calculated for SW13/cl.2 cells (D), A549 *Vim*^−/−^ cells (E) and MEF *Vim*^−/−^ (F). Results shown are average values ± SEM of 3 experiments totaling at least 14 cells per experimental condition. Statistical significance was evaluated with the Kruskal-Wallis plus Dunn’s test for multiple comparisons in (D), and by one-way ANOVA followed by Tukey’s test in (E, F). *p<0.05; **p<0.01; ***p<0.001; ****p<0.0001. In addition, a comparison between C328D and S49D,C328D in (F) was done with the Student-t test (#p<0.05).

Taken together, these results confirm the importance of the presence of C328 for vimentin droplet formation by diamide, and suggest that phosphorylation of vimentin at several sites, mainly at S49, could contribute to this effect, although is not sufficient to elicit droplet formation per se.

### Vimentin S49D,C328D forms droplets with the characteristics of biomolecular condensates

To explore the nature of vimentin S49D,C328D droplets, we assessed whether they shared characteristics of condensates. Therefore, we first studied the response of vimentin C328D assemblies and vimentin S49D,C328D droplets to incubation with the aliphatic alcohols 1,6-hexanediol, known to disperse condensates, and 2,5-hexanediol, which is ineffective and serves as a negative control [17]. Remarkably, vimentin C328D assemblies were not dispersed by either of the alcohols (Fig. 5A, left panels). However, pretreatment with diamide, which increases their circularity as illustrated in Fig. 2B, rendered them susceptible to dispersion by 1,6-hexanediol, but not 2,5-hexanediol (Fig. 5A, left panels). This indicates that diamide treatment induces phase separated biomolecular condensates of vimentin C328D, similar to its effect on vimentin wt [12]. Conversely, vimentin S49D,C328D assemblies were directly dispersed by 1,6-hexanediol, but not 2,5-hexanediol, consistent with their behavior as biomolecular condensates (Fig. 5A, right panels). Moreover, pretreatment with diamide did not significantly improve dispersion by 1,6-hexanediol or affected the lack of effect of 2,5-hexanediol. These effects were quantitated by determining the coefficient of variation of the fluorescence signal under the various treatments (Fig. 5B). Dispersion of the droplets, leading to a diffuse and more homogeneous signal resulted in a decrease of this parameter. This confirmed that vimentin C328D required pretreatment with diamide for dispersion, whereas vimentin S49D,C328D was readily dispersed by 1,6-hexanediol alone.

**Figure 5.**
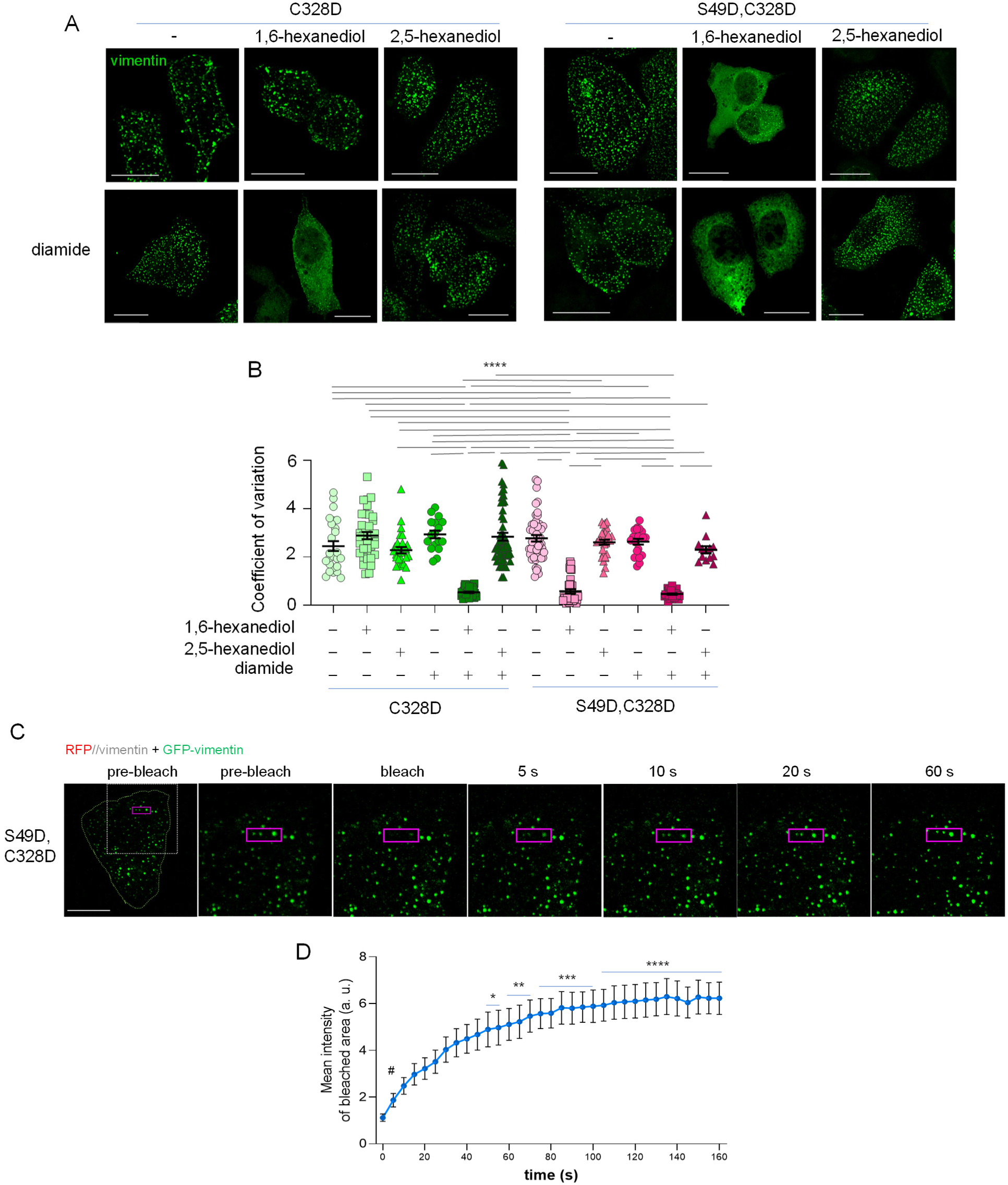
Vimentin S49D,C328D droplets share characteristics of biomolecular condensates. (A) Effect of aliphatic alcohols on the droplets formed by vimentin PTM mimetic mutants. SW13/cl.2 cells were transfected with vimentin C328D (left images) or S49D,C328D (right images), and incubated in serum free medium in the absence or presence of 1 mM diamide for 15 min, after which, 1,6-hexanediol or 2,5-hexanediol at 3.3% (w/v) final concentration, were added to the cells for 5 more min, as indicated. Cells were fixed and the vimentin constructs were monitored by immunofluorescence. (B) The coefficient of variation of the fluorescence intensity of the vimentin signal was monitored as an index of the dispersion of vimentin droplets, since remodeling of droplets into a diffuse background would cause a decrease in this parameter. Results are shown along the average values and SEM from at least 4 independent experiments totaling at least 13 cells per experimental condition. Statistical significance was analyzed with one-way ANOVA followed by the Tukey test for multiple comparisons. ****p<0.0001. (C) Vimentin S49D,C328D droplets undergo FRAP. SW13/cl.2 cells were cotransfected with plasmids RFP//vimentin S49D,C328D and GFP-vimentin S49D,C328D in a proportion 4:1, and 48 h later they were subjected to FRAP in a Leica SP5 microscope equipped with a thermostatized chamber, as detailed in methods. (D) Fluorescence intensity during recovery after photobleaching recorded by the FRAP wizard and the results from 14 assays are plotted as average values ± SEM. Statistical significance was evaluated by the Krukal-Wallis followed by the Dunn’s test (*p<0.05; **p<0.01; ***p<0.001; ****p<0.0001 vs postbleach 0 time. In addition, a comparison between the 0 and 5 min time points was done with the Student-t test (#p<0.001).

Phase separated condensates are dynamic compartments that exchange their content with the surroundings, which is usually explored by FRAP assays [22]. Therefore, we performed FRAP assays in cells transfected with a mixture of untagged and GFP-tagged vimentin constructs in a 4:1 proportion, in order to illuminate these structures while minimizing the perturbations elicited by the GFP-fusion protein (Fig. 5C) [12, 36]. We observed a fast recovery of fluorescence that could be detected at the first time point after bleaching (5 s) and progressed steadily. This confirmed the capability of vimentin S49D,C328D droplets of rapidly exchanging components with the surrounding environment (Fig. 5C and D).

In a previous work we showed that droplets formed by vimentin wt in cells treated with diamide displayed fast motility [12]. Nevertheless, diamide deeply alters the actin and tubulin cytoskeletons [24], for which motility of oxidant-elicited vimentin droplets may be conditioned by cytoskeletal disruption. In contrast, vimentin S49D,C328D should offer the opportunity to assess the behavior of vimentin phase separated condensates reminiscent of those formed under oxidative stress, in the absence of oxidant-elicited cytoskeletal disruption. The motion of vimentin S49D,C328D droplets in live cells adopted different patterns, with some droplets traveling variable distances and others remaining apparently static (Fig. 6A). The trajectories of several particles are illustrated in Fig. 6B, whereas the tracking of all particles in a single representative cell is shown in Fig. 6C. As it can be observed, movement directionality and length are highly variable. The speed and maximal distance traveled (distance between initial and final points) are quantitated in Fig. 6D and E. Remarkably, during monitorization of motility, several events suggestive of droplet fusion, i.e., coalescence of neighboring droplets, or splitting were observed (Fig. 6F).

**Figure 6.**
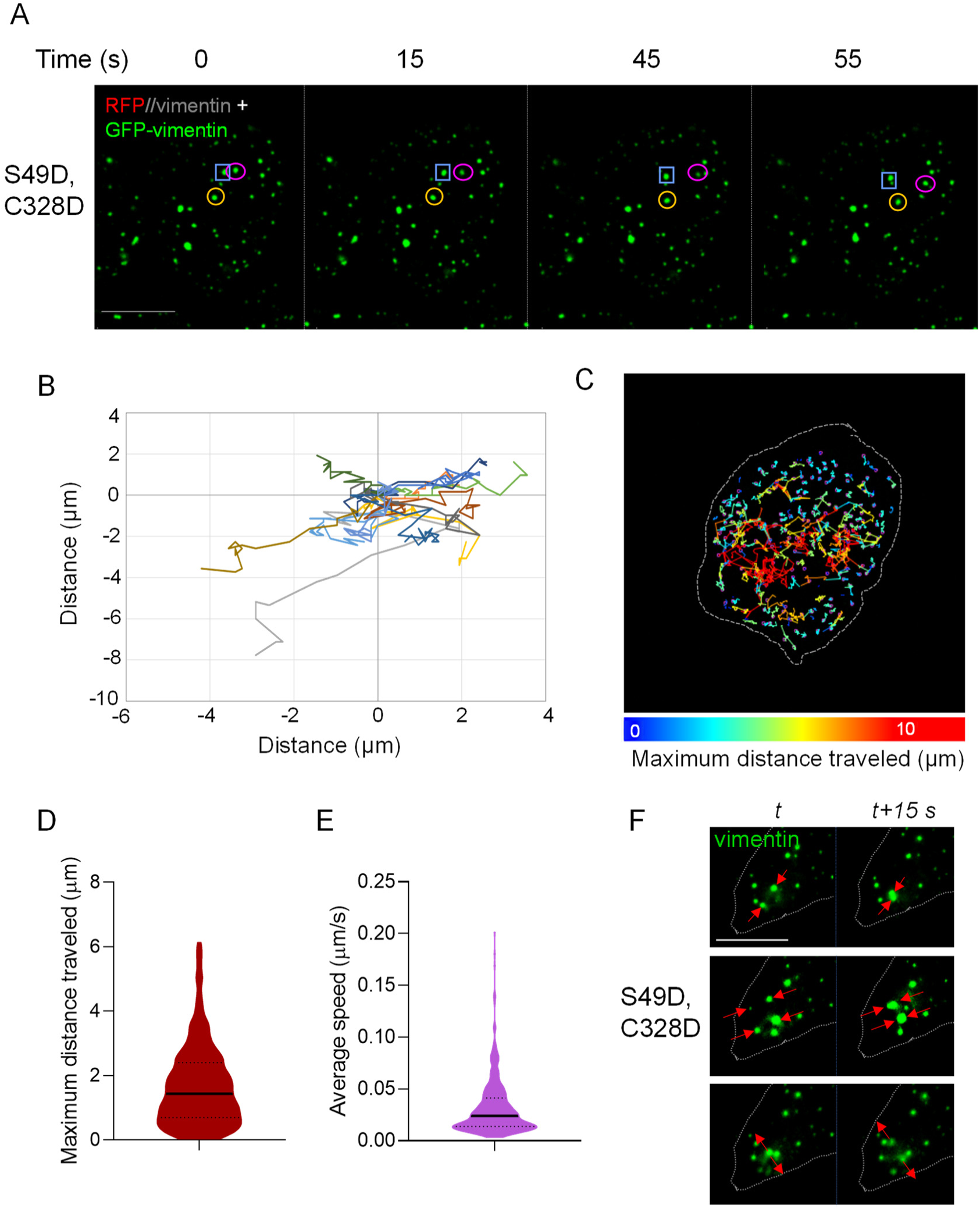
Motility of vimentin S49D,C328D droplets. (A) SW13/cl.2 cells were cotransfected with plasmids RFP//vimentin S49D,C328D and GFP-vimentin S49D,C328D in a proportion 4:1 and 48 h later, the motility of droplets was monitored in live cells. Three droplets are highlighted in different colors in order to illustrate their different positions along the time lapse. Scale bar, 20 µm. (B) The trajectories of 13 droplets were monitored and are shown in the diagram. (C) The movements of all droplets detected in one cell were monitored and are represented in different colors depending on the maximum distance traveled (given by a straight line between the initial and final positions) according to the color scale depicted at the bottom. The cell contour is marked by a dashed line. (D, E) The maximum distance traveled (D), and the maximum speed (E) of vimentin S49D,C328D droplets from three experiments monitoring more than 340 particles are represented. The solid line represents the median and the dotted lines, the 25 and 75 percentiles. (F) Representative images from the monitorization of vimentin S49D,C328D droplets motility showing events compatible with droplet coalescence (upper and middle sequences) or splitting (lower sequence). Scale bar, 10 µm.

### Effect of protein concentration on the formation of vimentin S49D,C328D condensates

Biomolecule concentration is an important determinant of condensate formation (reviewed in [22]). Here we observed that under normal cell culture conditions there was a positive correlation between the levels of vimentin S49D,C328D expressed in cells, estimated as the signal intensity after direct immunofluorescence, and the size of the condensates formed. This is illustrated in Fig. 7A, which shows a single field with cells spontaneously expressing different vimentin S49D,C328D levels as the result of transient transfection. Cells expressing low levels of vimentin S49D,C328D displayed fine dots, whereas those with high levels showed large droplets. The quantitation of these observations in individual cells shows a clear correlation between mean fluorescence intensity of the vimentin S49D,C328D signal and droplet size (Fig. 7B).

**Figure 7.**
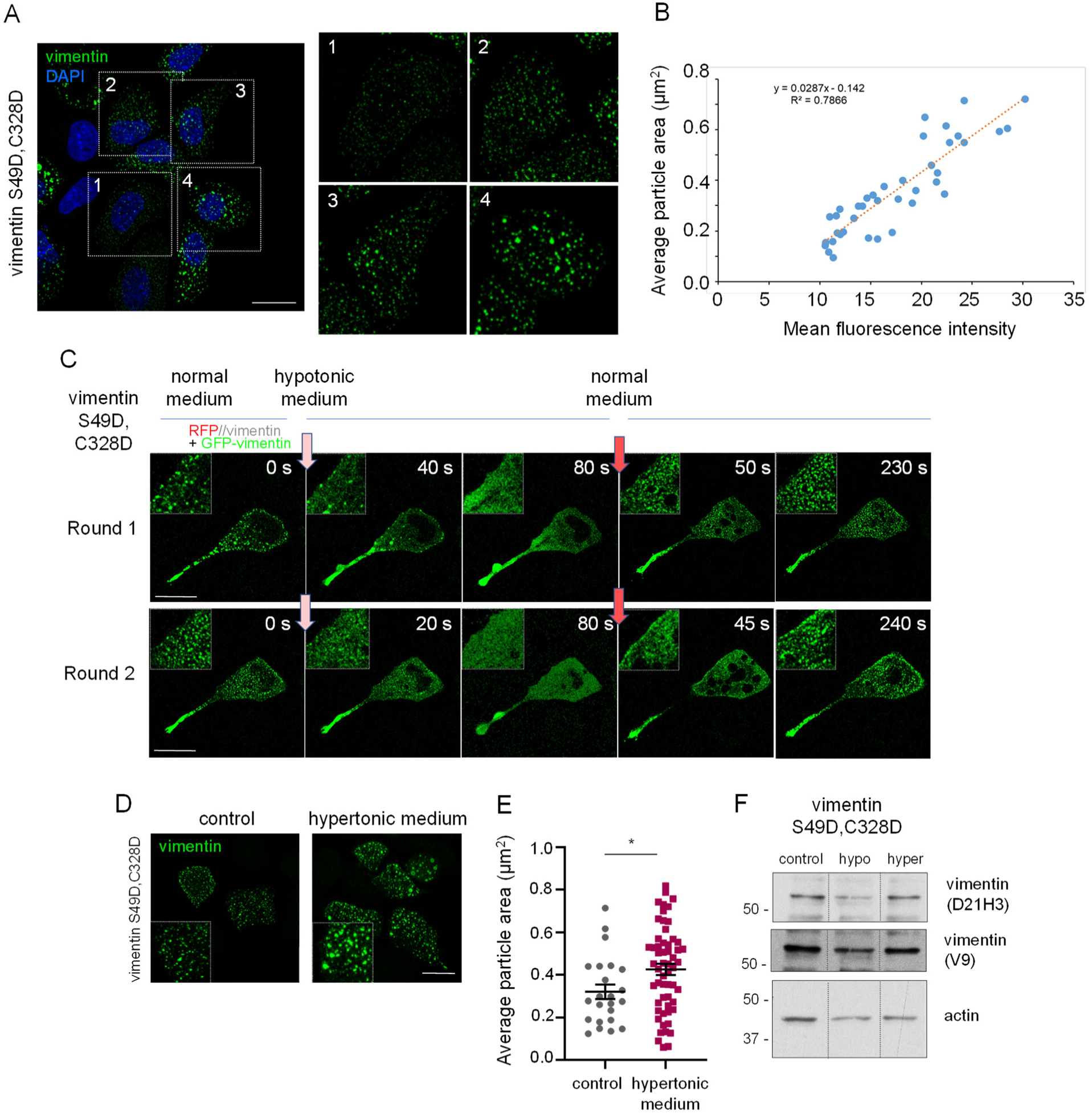
Effect of the cellular levels of vimentin S49D,C328D on the size of the assemblies formed. (A) Observation of the size distribution of vimentin S49D,C328D condensates depending on the level of the construct spontaneously achieved after transfection. SW13/cl.2 cells were transfected with the RFP//vimentin S49D,C328D plasmid and 48 h later the condensates were detected by immunofluorescence with the V9 anti-vimentin antibody. Images at the right depict enlarged views of cells expressing different levels of the vimentin S49D,C328D construct, according to the intensity of the fluorescent signal. (B) Fields with several transfected cells were randomly selected and the average particle area and mean fluorescence intensity of individual cells were measured. The graph shows the linear regression plot of data obtained from 39 cells from 4 independent experiments. (C) Effect of hypotonic shock on vimentin S49D,C328D assemblies. SW13/cl.2 cells were transfected with a mixture of RFP//vimentin S49D,C328D and GFP-vimentin S49D,C328D plasmids in a proportion 4:1 and 48 h later the response to changes in the osmolarity of the medium was monitored in live cells by fluorescence microscopy. Arrows indicate the moments at which the medium was changed to hypotonic (light pink) or isotonic (pink), and represent the time 0 for image acquisition under the corresponding experimental conditions. Images show the dispersion and reassembly of condensates in a single cell subjected to two rounds of hypotonic shock and recovery. Insets display enlarged areas of interest. Results shown are representative from the monitorization of at least 20 cells from 4 experiments. (D) Cells transfected as in (C) were subjected to hypertonic shock by incubation with culture medium supplemented with additional 150 mM NaCl for 5 min, after which, cells were imaged and the average particle area was monitored in a total of 23 and 57 cells for control and hypertonic shock cells, respectively, from 3 experiments. (E) The graph shows average values ± SEM. *p<0.05 by Student’s t-test. (F) Lysates from cells subjected to hypotonic or hypertonic shock for 5 min were analyzed by western blot with antibodies against vimentin N-terminus (D21H3) or C-terminus (V9). Levels of actin are shown as control. Dotted lines indicate where lanes from the same gel have been cropped.

A strategy to alter the concentration of the cellular contents is to subject cells to osmotic changes. Incubation in hypotonic medium would increase cell volume and dilute its contents, potentially dissociating condensates, whereas hypertonic medium would decrease cell volume resulting in higher biomolecule concentration and possibly increased condensate formation [53]. Here, we observed that short term incubation of cells expressing vimentin S49D,C328D in hypotonic medium led to the complete dispersion of condensates and appearance of the protein as a uniform diffuse background, indicative of solubilization (Fig. 7C, upper row). Notably, restituting normal medium restored phase separation with the rapid reappearance of droplets, confirming the reversibility of this process. Furthermore, the cycle of condensate dispersion and recovery could be repeated by switching the medium between normal and hypotonic (Fig. 7C, lower row). These transitions occurred in the time scale of seconds, illustrating the dynamicity of vimentin S49D,C328D structures. Conversely, incubation in hypertonic medium resulted in the appearance of larger droplets (Fig. 7D and E). As hypotonic shock has been reported to elicit vimentin degradation [54], we assessed the integrity of the vimentin S49D,C328D construct under these experimental conditions by western blot (Fig. 7F). Neither hypotonic nor hypertonic shock elicited changes in the mobility of the vimentin band with respect to the control condition, as detected with antibodies recognizing epitopes located in the N-terminal (D21H3) or C-terminal domains (V9), suggesting that loss of condensates was due to dispersion of the intact protein.

### Behavior of vimentin S49D,C328D in vitro

Given the capability of vimentin S49D,C328D to form biomolecular condensates in cells, we next explored its properties in vitro. We first assessed the capacity of this mutant to form filaments upon addition of NaCl, compared to vimentin wt, under standard polymerization conditions (Fig. 8A). Incubation of vimentin wt at 37°C for 1 h in hypotonic buffer (pH 7.0) led to the appearance of multiple fibrils and short filaments, whereas incubation in the presence of NaCl elicited the formation of characteristic long and well defined filaments [37] (Fig. 8A, upper images). In contrast, vimentin S49D,C328D formed short irregular structures in hypotonic buffer and larger but highly heterogeneous assemblies in the presence of NaCl (Fig. 8A, lower images). These included filamentous formations wider and shorter than vimentin wt filaments, and aggregates, which appeared in various proportions and combinations. The width of vimentin wt filaments and vimentin S49D,C328D assemblies under polymerization conditions is quantitated in Fig. 8B.

**Figure 8.**
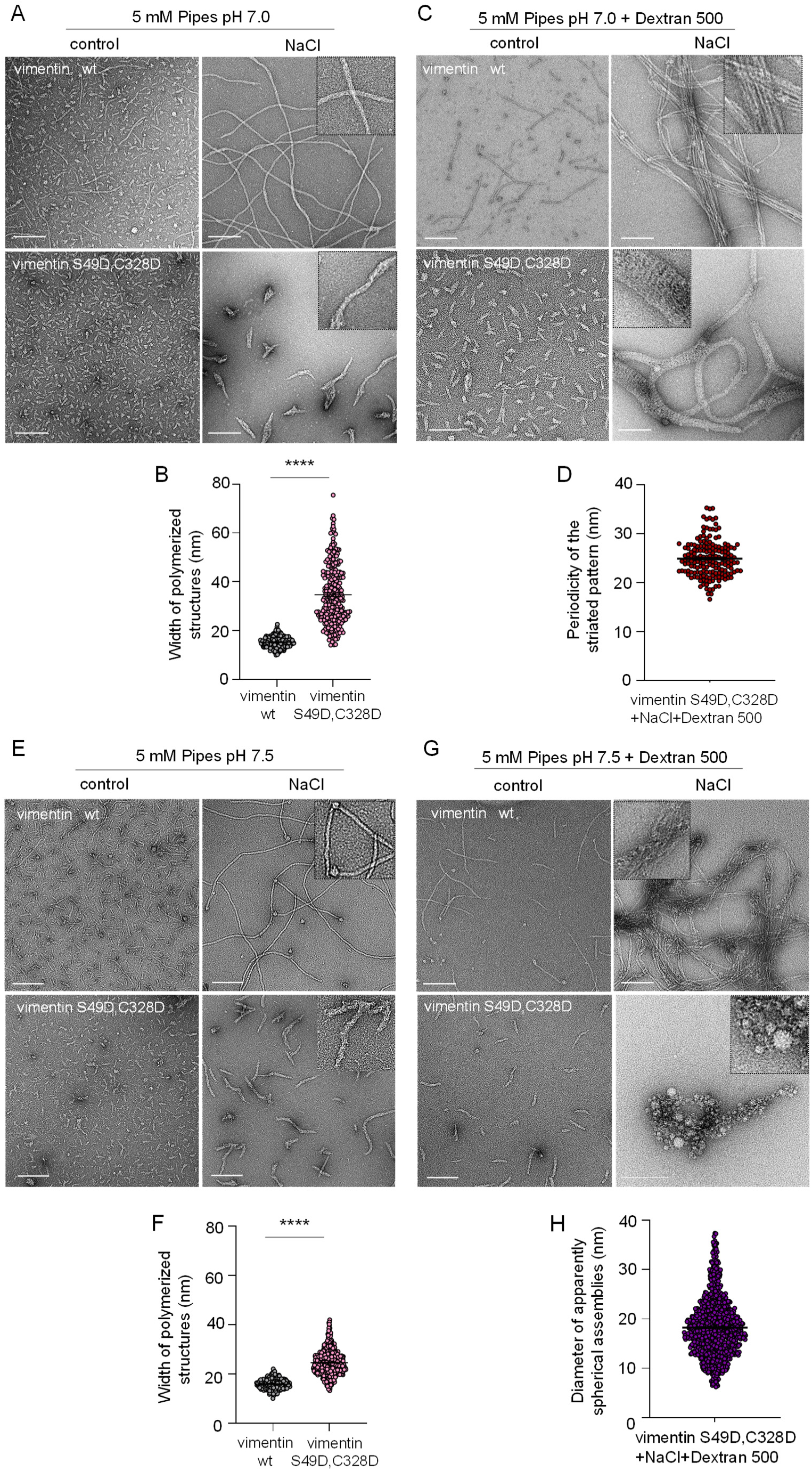
Vimentin S49D, C328D forms polymorphic assemblies in vitro in a pH and agglomeration dependent manner. (A) Purified vimentin wt or S49D, C328D was incubated at 37°C for 1 h in buffer at pH 7.0, in the absence or presence of 150 mM NaCl, as indicated. Incubation mixtures were fixed with glutaraldehyde and vimentin assemblies were visualized by TEM, as specified in Methods. The width of vimentin wt filaments and vimentin S49D,C328D assemblies is represented in (B). A total of 339 and 272 structures, respectively, were monitored from three independent assays. Statistical differences were evaluated with the Mann-Whitney test. ****p<0.0001 (C) The polymerization assay was performed in buffer at pH 7.0 in the presence of dextran 500. (D) Graph showing the distribution of the width of the axial repeat in the striated pattern of vimentin S49D,C328D paracrystals, totaling 174 measurements from at least five independent assays. (E) Vimentin wt or S49D,C328D was incubated at 37°C for 1 h in buffer at pH 7.5, in the absence or presence of 150 mM NaCl, as indicated. (F) The graph shows the width of the corresponding polymerized structures (totaling 335 and 780 determinations for vimentin wt and S49D,C328D, respectively). ****p<0.0001 by Mann-Whitney test. (G) Polymerization was carried out at pH 7.5 in the presence of dextran 500. (H) The graph shows the size distribution of the beads observed for vimentin S49D,C328D polymerized at pH 7.5 in the presence of dextran 500. More than 1000 “beads” from at least 3 different experiments were measured, and the average ± SEM are shown. Images shown are representative of at least three assays per experimental condition. ROIs are depicted in insets. Scale bars, 200 nm.

Since vimentin S49D, C328D polymerized structures were highly irregular, we hypothesized that droplet formation could require factors or conditions present in the cellular environment. As shown above, protein concentration plays an important role in condensate formation in cells. In fact, the cell cytoplasm is a crowded milieu, with an estimated concentration of total macromolecules between 100-400 mg/ml [55, 56], of which proteins can reach 100 mg/ml [57]. In contrast, protein concentration in the in vitro polymerization assay is three orders of magnitude lower (0.2 mg/ml). Multiple entities contribute to the intracellular density, including proteins and abundant structures such as ribosomes. Various crowding agents, which occupy a large volume and force macromolecules to concentrate, are used to mimic this environment in vitro. Therefore, we explored the effect of the crowding agent dextran 500 on the morphology of vimentin wt and S49D,C328D assemblies (Fig. 8C). We observed that the presence of dextran decreased the background of small precursors and increased the size of both vimentin wt and S49D,C328D assemblies, even in hypotonic buffer, possibly resulting from increased association of oligomeric species due to volume exclusion by the crowder (Fig. 8C, left images). Remarkably, induction of vimentin wt polymerization by addition of NaCl in the presence of the crowder resulted in the formation of broad networks constituted by robust and rather straight bundles of well defined filaments, some of which comprising more than 20 filaments (Fig. 8C, upper right). In sharp contrast, vimentin S49D,C328D polymerized in the presence of dextran 500 formed broad ribbon-like assemblies (at places wider than 100 nm) where filaments could not be individually visualized, but seemed to coalesce at some points yielding large areas of non-fibrillar texture (Fig. 8C, lower right). Actually, these areas displayed a striated pattern, resembling the paracrystal-like structures previously observed for several intermediate filament proteins in native or mutant forms [58–63]. Remarkably, the axial periodicity of the striated pattern of vimentin S49D,C328D displayed an average of 24.92 ± 0.28 nm (mean ± SEM, quantitated in Fig. 8D), which is reminiscent of what has been reported for lamin B2 (∼24.5 nm [60]) and truncated GFAP (∼22 nm [63]).

We have previously observed that mildly acidic pHi is not permissive for the formation of vimentin wt condensates in response to diamide in cells [24]. Moreover, pH is an important determinant of the assembly of vimentin and other intermediate filaments in vitro [24, 60, 64]. Therefore, we next explored the assembly of vimentin wt and S49D,C328D at pH 7.5 (Fig. 8E-H). In the absence of crowders, vimentin wt formed long regular filaments (Fig. 8E, upper right), with an average width of 15.97 ± 0.39 nm (mean ± SEM, quantitated in Fig. 8F), frequently associated with spherical formations of approximately 25.63 ± 0.79 nm of diameter (average ± SEM), often located at filament ends. In turn, S49D,C328D polymerized into highly heterogeneous structures, including fibrils and broad and short irregular filamentous assemblies (Fig. 8E, lower right), approximately 24.79 ± 0.22 nm wide (mean ± SEM, quantitated in Fig. 8F). Notably, the surface of these assemblies displayed a rough appearance, with some fibrils protruding from them as “thorns”. Incubation in hypotonic buffer at pH 7.5 in the presence of dextran 500 decreased the abundance of small particles and increased association of both vimentin wt and S49D,C328D into more filamentous structures. In turn, including dextran 500 in the polymerization mixture at pH 7.5 resulted in striking differences between vimentin wt and S49D,S328D (Fig. 8G). Vimentin wt polymerized into bundles that appeared curvier and more entangled than those formed at pH 7.0, displaying some poorly defined regions together with zones in which individual filaments could still be distinguished (Fig. 8G, upper right). In turn, polymerization of vimentin S49D,C328D in the presence of dextran at pH 7.5 resulted in the formation of clusters of protein assemblies of nearly spherical or beaded appearance, in which filamentous structures were not detected (Fig. 8G, lower right). The size of the “beads” was highly varied, and ranged from ∼10 to over 30 nm of diameter, with an average of 18.27 ± 0.18 nm (mean ± SEM, Fig. 8H). Nevertheless, these beads seemed interconnected, sometimes forming networks that could span several hundred nanometers.

Together, these results illustrate that variations in pH and in the agglomeration state elicit remarkable changes in the morphology of vimentin wt assemblies in vitro. Moreover, they also show that introducing the two PTM mimetic mutations S49D plus C328D, drastically impacts the organization of vimentin during polymerization. Therefore, the S49D,C328D mutations drive vimentin condensate formation in cells as well as the polymerization of the protein into paracrystal-like or beaded arrays in vitro.

## Discussion

We recently postulated that oxidative modifications could play a role in vimentin phase separation [22]. Here we have confirmed that the single cysteine residue of vimentin, C328, is substantially modified upon induction of cellular oxidative stress by treatment with diamide. However, under oxidative stress, vimentin can undergo numerous PTMs, including phosphorylation and various types of lipoxidation, not only at C328 but at many other residues [27]. Now, we have explored the role of some of these PTMs in condensate formation. Moreover, we have aimed to identify the minimum requirements to drive vimentin phase separation by introducing PTM mimetic mutations. Our results indicate that mimicking an oxidative modification of C328 and one phosphorylation at a precise site in the head domain is sufficient to elicit vimentin phase separation in cells. This implies introducing two negative charges at critical positions in the vimentin monomer, of which, perturbation of the 328 site is required for the phosphomimetic mutation to have an effect. These results highlight the importance of vimentin C328 in oxidant-elicited remodeling and establish a hierarchical relationship between cysteine modification and a minimal phosphorylation for phase separation.

Redox status has been shown to modulate the phase separation of various proteins. In particular, ApoE has been proposed to undergo cysteine oxidation mediated phase separation [65]. In turn, glutathionylation regulates the phase separation of RNA binding proteins [7], whereas the balance between cysteine modifications, persulfidation and sulfonylation, can modulate the transition between condensates and aggregates in models of aging [11]. In plants, cysteine residues of the redox sensor Radical-induced Cell Death1 (RCD1), regulate the formation of phase separated condensates [66], whereas condensates of the yeast ataxin-2 protein are disassembled by H_2_O_2_-elicited methionine oxidation [67].

The type III intermediate filaments, vimentin, GFAP and desmin are emerging as important players in the response to oxidative stress [14]. They possess a single conserved cysteine residue that is a hot spot for PTMs and is key for their remodeling upon exposure to oxidants and electrophiles [30]. We have recently shown that several oxidants elicit the remodeling of these intermediate filaments into biomolecular condensates [12]. The factors driving biomolecule phase separation and condensate formation are not completely understood, but include the formation of multivalent interactions between intrinsically disordered protein domains. PTMs can be critical modulators of these interactions and therefore are considered determinant for condensate formation [10].

Vimentin C328 can be the target of a great variety of modifications under oxidative stress, including mono, di and trioxidations, to sulfenic, sulfinic and sulfonic acids, respectively. In particular, the oxidant diamide has been found to elicit vimentin-vimentin disulfide formation, glutathionylation, and CoAlation [41, 43, 68, 69]. Here, we chose to mimic sulfinylation because the gain of a negative charge can be approximated through a cysteine to aspartate mutation. This mutation had a strong impact on vimentin assembly in cells, precluding the formation of filaments and leading to short irregular structures. In contrast, other mutations introduced at this site, such as substitutions by serine, alanine, histidine or even tryptophan, are relatively well tolerated and even confer resistance to disruption by oxidants [24, 31, 37]. A reason for the deleterious effect of the aspartate substitution may be the fact that in the mature filament, four cysteine residues from two adjacent tetramers appear to fall within close distance [30, 32], and therefore, introduction of negative charges at these sites may be disruptive. However, the C328D mutant did not completely recapitulate the impact of diamide treatment on cellular vimentin wt, as it was still susceptible to remodeling into more circular and numerous droplets upon treatment with the oxidant. Although many cellular mechanisms could contribute to this effect, we first explored whether modulating fast and reversible PTMs that occur on vimentin itself, namely glycosylation and phosphorylation, could influence droplet formation.

Vimentin is a hub for PTMs, which are in great part responsible for finely modulating its assembly. Site-specific O-GlcNAcylation has been reported to regulate the vimentin cytoskeleton. Several O-glycosylation sites have been postulated or identified, including, S7, T33, S34, S39, S49 and S55 [49, 70–72]. In particular, glycosylation of S49 has been proposed to play an important role in the homotypic assembly of vimentin into mature filaments in cells [49]. O-GlcNAcylation, has been reported to inhibit phase separation of proteins. Glycosylation of the brain GTPase activating protein SynGAP negatively regulates phase separation of the PSD-95/SynGAP complex in a site dependent manner [73, 74]. In turn, O-GlcNAc modifications on the RNA binding proteins YTHDF1 and YTHDF3 regulate the dynamics of stress granules during recovery from stress [75]. Our results with the deglycosylation inhibitor Thiamet G, also support that O-GlcNAcylation, either directly or indirectly, plays a negative role in vimentin phase separation. In turn, the vimentin sequence possesses at least 74 phosphorylation sites (according to www.phosphosite.org), and their coordinated modification is known to finely tune vimentin assembly in essential processes such as cell migration and division. Phosphorylation has been shown to rapidly modulate condensate formation. However, the nature of the effect, promoting or suppressing phase separation, is highly dependent on the particular protein and its context. Phosphorylation suppresses phase separation of the RNA binding protein FUS [76], whereas it may favor Tau phase separation [77]. In fact, within the same protein, the effect of phosphorylation on phase separation may depend on the site or the extent of the modification [78]. In this context, we have previously reported that, whereas diamide elicits vimentin droplets, its combination with the phosphatase inhibitor calyculin A, which induces vimentin hyperphosphorylation, results in the dispersion of these dots leading to diffuse vimentin [37]. Here, we have observed that the kinase inhibitors H-89 and staurosporine attenuate diamide-elicited C328D droplet formation, an effect that was not shared by the RhoK inhibitor Y27632, suggesting that RhoK target residues may be less relevant for this process. Because phosphorylation and glycosylation can occur on the same residues [72, 79], the possibility of interplay between these two PTMs in the modulation of phase separation can be proposed.

Therefore, both the evidence available in the literature and our own results point towards the potential involvement of deglycosylation and/or phosphorylation in vimentin disassembly and phase separation. The typical small assemblies formed by vimentin C328D evolved towards interconnected squiggles in the presence of inhibitors of these processes. We reasoned that if these effects were due to the direct modulation of vimentin PTMs, phosphomimetic vimentin mutants should have the opposite effect. Indeed, generation of various phosphomimetic mutants evidenced that, if combined with the C328D mutation, the S49D mutation elicited particle rounding more effectively and consistently across several cell types than other phosphomimetic point mutations. The S49 residue is especially interesting because it can be either phosphorylated or O-NAcglycosylated, two modifications with opposite repercussions on the stability of vimentin filaments. Whereas phosphorylation can contribute to filament disassembly, glycosylation has been shown to increase filament stability. Therefore, introducing a phosphomimetic residue at the S49 site could have the double functional impact of precluding a stabilizing modification and introducing a destabilizing one.

We observed that droplets formed by the double mutant vimentin S49D,C328D shared the typical features of biomolecular condensates. Therefore, in the case of vimentin, introduction of only two PTM mimics, one of them mimicking oxidized C328, switches vimentin organization from robust filaments into biomolecular condensates. This highlights the importance of C328, which, until now, was mainly supported by its requirement for vimentin efficient assembly and remodeling by oxidants [36, 37]. However, the results herein described provide evidence of the positive involvement of perturbations at this site in phase separation.

Nevertheless, it needs to be considered that condensates formed by vimentin S49D, C328D may differ from those formed by vimentin wt under oxidative stress. The double mutation studied here mimics just one of many PTM combinations potentially driving vimentin phase separation in cells. In contrast, it is likely that under oxidative stress, many vimentin proteoforms are generated, with different PTM combinations, that could display different propensity to phase separate. Therefore, the composition of vimentin condensates in oxidant treated cells could be highly heterogeneous, both with respect to the integrating vimentin proteoforms, as well as to the presence of other cellular components. This complexity further increases if we consider that perturbations at any given site in a protein sequence, either by mutation or PTM, can change the landscape or susceptibility to other PTMs.

On the other hand, the fact that vimentin S49D,C328D undergoes phase separation suggests that vimentin can drive condensate formation per se, more than just being a client protein. Moreover, this construct provides the advantage that condensates are stable enough to perform several studies, and do not require to expose cells to oxidative stress, which would be deleterious for many cellular structures in the long term. For instance, condensates formed by vimentin S49D,C328D can be monitored in time lapse experiments to assess their reversibility or dynamics during the course of minutes to hours. In particular, dilution of cellular contents through hypotonic shock readily disperses vimentin S49D,C328D condensates, which reform when osmotic conditions are restored.

Similarly to the results obtained in cells, the two PTM mimetic mutations also severely impaired filament assembly in vitro, although the structures formed by vimentin S49D,C328D under standard polymerization conditions were highly heterogeneous and did not resemble droplets. Remarkably, incorporation of dextran 500 in vimentin incubation mixtures increased the abundance of filamentous or polymerized structures, even in hypotonic medium, probably due the concentration of the protein in the volume that remains hydrodynamically available, or “free” volume, that is estimated to be a 30 to 45% of the total volume of the solution [80]. In turn, polymerization in the presence of the crowding agent elicited the formation of robust bundles of vimentin wt, which formed an interconnected network on the grid. This is in sharp contrast with the typical individual filaments formed under dilute conditions. Since vimentin bundles have often been observed by electron microscopy in cells [23], it could be speculated that the crowded cellular environment would favor vimentin polymerization and bundling. Under the same polymerization conditions, vimentin S49D,C328D formed assemblies reminiscent of paracrystals, with an striated pattern of an average periodicity of approximately 25 nm. Interestingly, paracrystals of other intermediate filaments have been reported, including neurofilaments, lamins and GFAP fragments, which display similar periodicity [58–63]. The appearance of this striated pattern is due to the highly ordered periodic assembly of the proteins and the greater protein density in the overlapping regions, which retain less contrasting agent [63]. Although formation of paracrystals of other intermediate filament proteins has been reported to require divalent cations, this requirement appears not to be present in the case of vimentin S49D,C328D. The precise disposition of vimentin S49D,C328D in the paracrystals is not known, but it could represent a special alignment of the rods of the tetramers.

Remarkably, rising the pH of the incubation from 7.0 to 7.5 had a marked impact on the assembly of vimentin wt, and even a more profound effect on the vimentin S49D,C328D mutant. At pH 7.5, vimentin wt polymerized into filaments associated with round assemblies, similar to those previously observed at pH 8.0 in diluted conditions [24], and into entangled and poorly defined bundles in the presence of dextran 500. Under these conditions, the pattern of polymerized vimentin S49D,C328D was particularly striking, apparently forming “beaded” arrays or clusters of spherical densities that could form large branched collections. The average diameter of the beads was close to 20 nm. This raises the possibility that they reflect a particular arrangement of protein domains. A beaded pattern has been previously observed by atomic force microscopy of vimentin [81], and interpreted in terms of the different protein densities of ULF segments, depending on the alignment of vimentin tetramers. Also, a beaded pattern has been appreciated in studies of keratins [82]. On the other hand, certain desmin mutants have been shown to form round aggregates of approximately 30 nm in diameter [83, 84]. Additional work would be needed to gain insight into the nature of vimentin S49D,C328D formations in vitro, as well as to identify potential conditions and/or components that may drive droplet formation.

In summary, the results with vimentin wt indicate a regulation of condensate formation by the interplay of PTMs in cells, which can be recapitulated by a minimum number of mutations mimicking C328 oxidation and one site-specific phosphorylation. Moreover, our observations illustrate that macromolecular agglomeration and pH variations in the physiological range modulate of vimentin wt and S49D,C328D polymerization in vitro. These effects, which could be further regulated by additional factors, such as presence of divalent cations, or the interplay between PTM, could have an important counterpart in vimentin organization in cells.

## Acknowledgements

We thank Dr. Silvia Zorrilla and Dr. Miguel Ángel Robles Ramos, from CIB Margarita Salas, CSIC, Spain, for helpful comments and discussion.

## Abbreviations

FRAP: fluorescence recovery after photobleaching
GFP: green fluorescent protein
GlcNAc: N-acetylglucosamine
MALPEG: methoxypolyethylene glycol maleimide
MEF: murine embryonic fibroblast
PTM: posttranslational modification
ULF: unit length filament

## References

[1] S. Alberti, A.A. Hyman, Biomolecular condensates at the nexus of cellular stress, protein aggregation disease and ageing, Nat Rev Mol Cell Biol 22(3) (2021) 196–213.

[2] S. Alberti, Phase separation in biology, Curr Biol 27(20) (2017) R1097–R1102.

[3] S.F. Banani, H.O. Lee, A.A. Hyman, M.K. Rosen, Biomolecular condensates: organizers of cellular biochemistry, Nat Rev Mol Cell Biol 18(5) (2017) 285–298.

[4] J.B. Woodruff, B. Ferreira Gomes, P.O. Widlund, J. Mahamid, A. Honigmann, A.A. Hyman, The Centrosome Is a Selective Condensate that Nucleates Microtubules by Concentrating Tubulin, Cell 169(6) (2017) 1066–1077 e10.

[5] J.B. Woodruff, A.A. Hyman, E. Boke, Organization and Function of Non-dynamic Biomolecular Condensates, Trends Biochem Sci 43(2) (2018) 81–94.

[6] S. Alberti, D. Dormann, Liquid-Liquid Phase Separation in Disease, Annu Rev Genet 53 (2019) 171–194.

[7] H.J. Choi, J.Y. Lee, K. Kim, Glutathionylation on RNA-binding proteins: a regulator of liquid‒liquid phase separation in the pathogenesis of amyotrophic lateral sclerosis, Experimental & molecular medicine 55(4) (2023) 735–744.

[8] G. Rivas, A.P. Minton, Influence of Nonspecific Interactions on Protein Associations: Implications for Biochemistry In Vivo, Annu Rev Biochem 91 (2022) 321–351.

[9] G. Rivas, A.P. Minton, Surfaces as frameworks for intracellular organization, Trends Biochem Sci 49(11) (2024) 942–954.

[10] W.T. Snead, A.S. Gladfelter, The Control Centers of Biomolecular Phase Separation: How Membrane Surfaces, PTMs, and Active Processes Regulate Condensation, Mol Cell 76(2) (2019) 295–305.

[11] T. Vignane, M. Hugo, C. Hoffmann, A. Katsouda, J. Petric, H. Wang, M. Miler, F. Comas, D. Petrovic, S. Chen, J.L. Miljkovic, J.L. Morris, S.R. Chowdhury, J. Prudent, N. Polovic, M.P. Murphy, A. Papapetropoulos, D. Milovanovic, M.R. Filipovic, Protein thiol alterations drive aberrant phase separation in aging, bioRxiv (2023) 10.1101/2023.11.07.566021.

[12] P. Martínez-Cenalmor, A.E. Martínez, D. Moneo-Corcuera, P. González-Jiménez, D. Pérez-Sala, Oxidative stress elicits the remodeling of vimentin filaments into biomolecular condensates, Redox biology 75 (2024) 103282.

[13] J.E. Eriksson, T. Dechat, B. Grin, B. Helfand, M. Mendez, H.M. Pallari, R.D. Goldman, Introducing intermediate filaments: from discovery to disease, J Clin Invest 119(7) (2009) 1763–71.

[14] D. Pérez-Sala, R. Quinlan, The redox-responsive roles of intermediate filaments in cellular stress detection, integration and mitigation, Curr Opin Cell Biol 86 (2024) 102283.

[15] N.T. Snider, M.B. Omary, Post-translational modifications of intermediate filament proteins: mechanisms and functions, Nat Rev Mol Cell Biol 15(3) (2014) 163–77.

[16] H. Herrmann, M. Haner, M. Brettel, S.A. Muller, K.N. Goldie, B. Fedtke, A. Lustig, W.W. Franke, U. Aebi, Structure and assembly properties of the intermediate filament protein vimentin: the role of its head, rod and tail domains, J Mol Biol 264(5) (1996) 933–53.

[17] Y. Lin, E. Mori, M. Kato, S. Xiang, L. Wu, I. Kwon, S.L. McKnight, Toxic PR Poly-Dipeptides Encoded by the C9orf72 Repeat Expansion Target LC Domain Polymers, Cell 167(3) (2016) 789–802 e12.

[18] X. Zhou, Y. Lin, M. Kato, E. Mori, G. Liszczak, L. Sutherland, V.O. Sysoev, D.T. Murray, R. Tycko, S.L. McKnight, Transiently structured head domains control intermediate filament assembly, Proc Natl Acad Sci U S A 118(8) (2021) e2022121118.

[19] X. Zhou, M. Kato, S.L. McKnight, How do disordered head domains assist in the assembly of intermediate filaments?, Curr Opin Cell Biol 85 (2023) 102262.

[20] K.M. Ridge, J.E. Eriksson, M. Pekny, R.D. Goldman, Roles of vimentin in health and disease, Genes Dev 36(7-8) (2022) 391–407.

[21] I. Ramos, K. Stamatakis, C.L. Oeste, D. Perez-Sala, Vimentin as a Multifaceted Player and Potential Therapeutic Target in Viral Infections, International journal of molecular sciences 21(13) (2020) 4675.

[22] D. Pérez-Sala, S. Zorrilla, Versatility of vimentin assemblies: From filaments to biomolecular condensates and back, Eur J Cell Biol 104(2) (2025) 151487.

[23] B. Renganathan, S.A. Adam, V.I. Gelfand, Vimentin Intermediate Filaments: A Paradigm Shift From Static Structure to Dynamic Cytoplasmic Network, Bioessays 48(3) (2026) e70125.

[24] A.E. Martínez, P. Martínez-Cenalmor, P. González-Jiménez, C. Vidal-Verdú, M.A. Pajares, D. Pérez-Sala, Vimentin remodeling in response to oxidants and electrophiles is modulated by pH, Sci Adv 12 (2026) eaed9655.

[25] J.E. Eriksson, T. He, A.V. Trejo-Skalli, A.-S. Härmälä-Braskén, J. Hellman, Y.-H. Chou, R.D. Goldman, Specific in vivo phosphorylation sites determine the assembly dynamics of vimentin intermediate filaments, J Cell Sci 117 (2004) 919–932.

[26] M. Matsuyama, H. Tanaka, A. Inoko, H. Goto, S. Yonemura, K. Kobori, Y. Hayashi, E. Kondo, S. Itohara, I. Izawa, M. Inagaki, Defect of mitotic vimentin phosphorylation causes microophthalmia and cataract via aneuploidy and senescence in lens epithelial cells, J Biol Chem 288(50) (2013) 35626–35.

[27] E. Griesser, V. Vemula, A. Mónico, D. Pérez-Sala, M. Fedorova, Dynamic posttranslational modifications of cytoskeletal proteins unveil hot spots under nitroxidative stress, Redox biology 44 (2021) 102014.

[28] P.V. Hornbeck, J.M. Kornhauser, S. Tkachev, B. Zhang, E. Skrzypek, B. Murray, V. Latham, M. Sullivan, PhosphoSitePlus: a comprehensive resource for investigating the structure and function of experimentally determined post-translational modifications in man and mouse, Nucleic Acids Res 40(Database issue) (2012) D261–70.

[29] A. Viedma-Poyatos, M.A. Pajares, D. Pérez-Sala, Type III intermediate filaments as targets and effectors of electrophiles and oxidants, Redox biology 36 (2020) 101582.

[30] M.A. Pajares, D. Pérez-Sala, Type III intermediate filaments in redox interplay: key role of the conserved cysteine residue, Biochem Soc Trans 52 (2024) 849–860.

[31] P. González-Jiménez, S. Duarte, A. Martínez-Fernández, E. Navarro-Carrasco, V. Lalioti, M.A. Pajares, D. Pérez-Sala, Vimentin single cysteine residue acts as a tunable sensor for network organization and as a key for actin remodeling in response to oxidants and electrophiles, Redox biology 64 (2023) 102756.

[32] M. Eibauer, M.S. Weber, R. Kronenberg-Tenga, C.T. Beales, R. Boujemaa-Paterski, Y. Turgay, S. Sivagurunathan, J. Kraxner, S. Koster, R.D. Goldman, O. Medalia, Vimentin filaments integrate low-complexity domains in a complex helical structure, Nat Struct Mol Biol 31 (2024) 939–949.

[33] B.T. Helfand, M.G. Mendez, S.N. Murthy, D.K. Shumaker, B. Grin, S. Mahammad, U. Aebi, T. Wedig, Y.I. Wu, K.M. Hahn, M. Inagaki, H. Herrmann, R.D. Goldman, Vimentin organization modulates the formation of lamellipodia, Mol Biol Cell 22(8) (2011) 1274–89.

[34] M.P. Serres, M. Samwer, B.A. Truong Quang, G. Lavoie, U. Perera, D. Gorlich, G. Charras, M. Petronczki, P.P. Roux, E.K. Paluch, F-Actin Interactome Reveals Vimentin as a Key Regulator of Actin Organization and Cell Mechanics in Mitosis, Dev Cell 52(2) (2020) 210–222 e7.

[35] V. Lalioti, D. Moneo-Corcuera, D. Pérez-Sala, Key role of vimentin in the organization of the primary cilium, bioRxiv (2024) 10.1101/2024.01.17.576004.

[36] D. Pérez-Sala, C.L. Oeste, A.E. Martínez, B. Garzón, M.J. Carrasco, F.J. Cañada, Vimentin filament organization and stress sensing depend on its single cysteine residue and zinc binding, Nature communications 6 (2015) 7287.

[37] A. Mónico, S. Duarte, M.A. Pajares, D. Pérez-Sala, Vimentin disruption by lipoxidation and electrophiles: role of the cysteine residue and filament dynamics, Redox biology 23 (2019) 101098.

[38] A. Basu, T. Krug, B. du Pont, Q. Huang, S. Sun, S.A. Adam, R.D. Goldman, D.A. Weitz, Vimentin undergoes liquid-liquid phase separation to form droplets which wet and stabilize actin fibers, Proc Natl Acad Sci U S A 122(10) (2025) e2418624122.

[39] A. Mónico, E. Rodríguez-Senra, F.J. Cañada, S. Zorrilla, D. Pérez-Sala, Drawbacks of dialysis procedures for removal of EDTA, PLoS ONE 12(1) (2017) e0169843.

[40] S. Duarte, T. Melo, R. Domingues, J.d.D. Alché, D. Pérez-Sala, Insight into the cellular effects of nitrated phospholipids: evidence for pleiotropic mechanisms of action, Free Rad Biol Med 144 (2019) 192–202.

[41] M. Fratelli, H. Demol, M. Puype, S. Casagrande, I. Eberini, M. Salmona, V. Bonetto, M. Mengozzi, F. Duffieux, E. Miclet, A. Bachi, J. Vandekerckhove, E. Gianazza, P. Ghezzi, Identification by redox proteomics of glutathionylated proteins in oxidatively stressed human T lymphocytes, Proc Natl Acad Sci U S A 99 (2002) 3505–3510.

[42] M. Kaus-Drobek, N. Mucke, R.H. Szczepanowski, T. Wedig, M. Czarnocki-Cieciura, M. Polakowska, H. Herrmann, A. Wyslouch-Cieszynska, M. Dadlez, Vimentin S-glutathionylation at Cys328 inhibits filament elongation and induces severing of mature filaments in vitro, FEBS J 287 (2020) 5304–5322.

[43] N. Goya-Iglesias, B. Yu, I. Gout, D. Pérez-Sala, Type III intermediate filaments as novel CoAlation targets, Redox Rep 31(1) (2026) 2692797.

[44] S. Akter, L. Fu, Y. Jung, M.L. Conte, J.R. Lawson, W.T. Lowther, R. Sun, K. Liu, J. Yang, K.S. Carroll, Chemical proteomics reveals new targets of cysteine sulfinic acid reductase, Nat Chem Biol 14(11) (2018) 995–1004.

[45] A. Mónico, J. Guzman-Caldentey, M.A. Pajares, S. Martin-Santamaria, D. Pérez-Sala, Molecular Insight into the Regulation of Vimentin by Cysteine Modifications and Zinc Binding, Antioxidants (Basel) 10(7) (2021) 1039.

[46] L. Makmura, M. Hamann, A. Areopagita, S. Furuta, A. Munoz, J. Momand, Development of a sensitive assay to detect reversibly oxidized protein cysteine sulfhydryl groups, Antioxid Redox Signal 3(6) (2001) 1105–18.

[47] Y. Miyata, J.N. Rauch, U.K. Jinwal, A.D. Thompson, S. Srinivasan, C.A. Dickey, J.E. Gestwicki, Cysteine reactivity distinguishes redox sensing by the heat-inducible and constitutive forms of heat shock protein 70, Chemistry & biology 19(11) (2012) 1391–9.

[48] M.A. Wilson, The role of cysteine oxidation in DJ-1 function and dysfunction, Antioxid Redox Signal 15(1) (2011) 111–22.

[49] H.J. Tarbet, L. Dolat, T.J. Smith, B.M. Condon, E.T. O’Brien, 3rd, R.H. Valdivia, M. Boyce, Site-specific glycosylation regulates the form and function of the intermediate filament cytoskeleton, eLife 7 (2018) e31807.

[50] D.M. Toivola, P. Strnad, A. Habtezion, M.B. Omary, Intermediate filaments take the heat as stress proteins, Trends Cell Biol 20(2) (2010) 79–91.

[51] S.A. Yuzwa, M.S. Macauley, J.E. Heinonen, X. Shan, R.J. Dennis, Y. He, G.E. Whitworth, K.A. Stubbs, E.J. McEachern, G.J. Davies, D.J. Vocadlo, A potent mechanism-inspired O-GlcNAcase inhibitor that blocks phosphorylation of tau in vivo, Nat Chem Biol 4(8) (2008) 483–90.

[52] S.Y. Kim, S.J. Jeong, J.H. Park, W. Cho, Y.H. Ahn, Y.H. Choi, G.T. Oh, R.L. Silverstein, Y.M. Park, Plasma Membrane Localization of CD36 Requires Vimentin Phosphorylation; A Mechanism by Which Macrophage Vimentin Promotes Atherosclerosis, Front Cardiovasc Med 9 (2022) 792717.

[53] P. Li, P. Chen, F. Qi, J. Shi, W. Zhu, J. Li, P. Zhang, H. Xie, L. Li, M. Lei, X. Ren, W. Wang, L. Zhang, X. Xiang, Y. Zhang, Z. Gao, X. Feng, W. Du, X. Liu, L. Xia, B.F. Liu, Y. Li, High-throughput and proteome-wide discovery of endogenous biomolecular condensates, Nat Chem 16(7) (2024) 1101–1112.

[54] L. Pan, P. Zhang, F. Hu, R. Yan, M. He, W. Li, J. Xu, K. Xu, Hypotonic Stress Induces Fast, Reversible Degradation of the Vimentin Cytoskeleton via Intracellular Calcium Release, Adv Sci (Weinh) 6(18) (2019) 1900865.

[55] G. Guigas, C. Kalla, M. Weiss, Probing the nanoscale viscoelasticity of intracellular fluids in living cells, Biophysical journal 93(1) (2007) 316–23.

[56] A.T. Molines, J. Lemiere, M. Gazzola, I.E. Steinmark, C.H. Edrington, C.T. Hsu, P. Real-Calderon, K. Suhling, G. Goshima, L.J. Holt, M. Thery, G.J. Brouhard, F. Chang, Physical properties of the cytoplasm modulate the rates of microtubule polymerization and depolymerization, Dev Cell 57(4) (2022) 466–479 e6.

[57] B.J. Zeskind, C.D. Jordan, W. Timp, L. Trapani, G. Waller, V. Horodincu, D.J. Ehrlich, P. Matsudaira, Nucleic acid and protein mass mapping by live-cell deep-ultraviolet microscopy, Nat Methods 4(7) (2007) 567–9.

[58] P. Traub, A. Scherbarth, J. Willingale-Theune, U. Traub, Large scale co-isolation of vimentin and nuclear lamins from ehrlich ascites tumor cells cultured in vitro, Prep Biochem 18(4) (1988) 381–404.

[59] R.D. Moir, R.A. Quinlan, M. Stewart, Expression and characterization of human lamin C, FEBS Lett 268(1) (1990) 301–5.

[60] E. Heitlinger, M. Peter, M. Haner, A. Lustig, U. Aebi, E.A. Nigg, Expression of chicken lamin B2 in Escherichia coli: characterization of its structure, assembly, and molecular interactions, J Cell Biol 113(3) (1991) 485–95.

[61] S. Heins, P.C. Wong, S. Muller, K. Goldie, D.W. Cleveland, U. Aebi, The rod domain of NF-L determines neurofilament architecture, whereas the end domains specify filament assembly and network formation, J Cell Biol 123(6 Pt 1) (1993) 1517–33.

[62] R.A. Quinlan, R.D. Moir, M. Stewart, Expression in Escherichia coli of fragments of glial fibrillary acidic protein: characterization, assembly properties and paracrystal formation, J Cell Sci 93 (Pt 1) (1989) 71–83.

[63] N.H. Lin, W.S. Jian, M.D. Perng, Deletions in Glial Fibrillary Acidic Protein Leading to Alterations in Intermediate Filament Assembly and Network Formation, International journal of molecular sciences 26(5) (2025) 1913.

[64] N. Mucke, T. Wedig, A. Burer, L.N. Marekov, P.M. Steinert, J. Langowski, U. Aebi, H. Herrmann, Molecular and biophysical characterization of assembly-starter units of human vimentin, J Mol Biol 340(1) (2004) 97–114.

[65] N. La Cunza, L.X. Tan, T. Thamban, C.J. Germer, G. Rathnasamy, K.A. Toops, A. Lakkaraju, Mitochondria-dependent phase separation of disease-relevant proteins drives pathological features of age-related macular degeneration, JCI insight 6(9) (2021) e142254.

[66] J. Xing, K. Li, W. Duan, Z. Duan, Y. Zhang, A. Guo, S. Wang, N. Ding, Y. Zou, X. Wei, X. Zhao, J. Zhang, Y. Song, H. Shi, S. Guo, C.P. Song, Phase separation of the redox sensor RCD1 mediates differential ROS signals to regulate plant growth and stress responses, Mol Plant 19(4) (2026) 868–886.

[67] M. Kato, Y.S. Yang, B.M. Sutter, Y. Wang, S.L. McKnight, B.P. Tu, Redox State Controls Phase Separation of the Yeast Ataxin-2 Protein via Reversible Oxidation of Its Methionine-Rich Low-Complexity Domain, Cell 177(3) (2019) 711–721 e8.

[68] D. Su, M.J. Gaffrey, J. Guo, K.E. Hatchell, R.K. Chu, T.R. Clauss, J.T. Aldrich, S. Wu, S. Purvine, D.G. Camp, R.D. Smith, B.D. Thrall, W.J. Qian, Proteomic identification and quantification of S-glutathionylation in mouse macrophages using resin-assisted enrichment and isobaric labeling, Free Radic Biol Med 67 (2014) 460–70.

[69] D.F. Vileigas, R.P. da Silva, B. Dempsey, M.P. Massafera, M.P. Pinz, F.C. Meotti, Redox proteomics workflow to unveil extracellular targets of oxidation in vascular endothelial cells, J Proteomics 321 (2025) 105506.

[70] C. Slawson, T. Lakshmanan, S. Knapp, G.W. Hart, A mitotic GlcNAcylation/phosphorylation signaling complex alters the posttranslational state of the cytoskeletal protein vimentin, Mol Biol Cell 19(10) (2008) 4130–40.

[71] Z. Wang, A. Pandey, G.W. Hart, Dynamic interplay between O-linked N-acetylglucosaminylation and glycogen synthase kinase-3-dependent phosphorylation, Mol Cell Proteomics 6(8) (2007) 1365–79.

[72] Z. Wang, N.D. Udeshi, C. Slawson, P.D. Compton, K. Sakabe, W.D. Cheung, J. Shabanowitz, D.F. Hunt, G.W. Hart, Extensive crosstalk between O-GlcNAcylation and phosphorylation regulates cytokinesis, Science signaling 3(104) (2010) ra2.

[73] P. Lv, Y. Du, C. He, L. Peng, X. Zhou, Y. Wan, M. Zeng, W. Zhou, P. Zou, C. Li, M. Zhang, S. Dong, X. Chen, O-GlcNAcylation modulates liquid-liquid phase separation of SynGAP/PSD-95, Nat Chem 14(7) (2022) 831–840.

[74] X. Li, L. Pinou, Y. Du, X. Chen, C. Liu, Emerging roles of O-glycosylation in regulating protein aggregation, phase separation, and functions, Current opinion in chemical biology 75 (2023) 102314.

[75] Y. Chen, R. Wan, Z. Zou, L. Lao, G. Shao, Y. Zheng, L. Tang, Y. Yuan, Y. Ge, C. He, S. Lin, O-GlcNAcylation determines the translational regulation and phase separation of YTHDF proteins, Nat Cell Biol 25(11) (2023) 1676–1690.

[76] Z. Monahan, V.H. Ryan, A.M. Janke, K.A. Burke, S.N. Rhoads, G.H. Zerze, R. O’Meally, G.L. Dignon, A.E. Conicella, W. Zheng, R.B. Best, R.N. Cole, J. Mittal, F. Shewmaker, N.L. Fawzi, Phosphorylation of the FUS low-complexity domain disrupts phase separation, aggregation, and toxicity, EMBO J 36(20) (2017) 2951–2967.

[77] S.F. Longfield, M. Mollazade, T.P. Wallis, R.S. Gormal, M. Joensuu, J.R. Wark, A.J. van Waardenberg, C. Small, M.E. Graham, F.A. Meunier, R. Martinez-Marmol, Tau forms synaptic nano-biomolecular condensates controlling the dynamic clustering of recycling synaptic vesicles, Nature communications 14(1) (2023) 7277.

[78] J. Chen, W. Ma, J. Yu, X. Wang, H. Qian, P. Li, H. Ye, Y. Han, Z. Su, M. Gao, Y. Huang, (-)-Epigallocatechin-3-gallate, a Polyphenol from Green Tea, Regulates the Liquid-Liquid Phase Separation of Alzheimer’s-Related Protein Tau, Journal of agricultural and food chemistry 71(4) (2023) 1982–1993.

[79] G.W. Hart, C. Slawson, G. Ramirez-Correa, O. Lagerlof, Cross talk between O-GlcNAcylation and phosphorylation: roles in signaling, transcription, and chronic disease, Annu Rev Biochem 80 (2011) 825–58.

[80] G. Migliorini, J.C. Vidos, J. Hamacek, A. Yethiraj, F. Piazza, Physical properties of dextran solutions as model crowding media, arXiv (2026) arXiv:2604.09373v1.

[81] S. Ando, K. Nakao, R. Gohara, Y. Takasaki, K. Suehiro, Y. Oishi, Morphological analysis of glutaraldehyde-fixed vimentin intermediate filaments and assembly-intermediates by atomic force microscopy, Biochim Biophys Acta 1702(1) (2004) 53–65.

[82] L. Milam, H.P. Erickson, Visualization of a 21-nm axial periodicity in shadowed keratin filaments and neurofilaments, J Cell Biol 94(3) (1982) 592–6.

[83] H. Bar, N. Mucke, A. Kostareva, G. Sjoberg, U. Aebi, H. Herrmann, Severe muscle disease-causing desmin mutations interfere with in vitro filament assembly at distinct stages, Proc Natl Acad Sci U S A 102(42) (2005) 15099–104.

[84] H. Herrmann, H. Bar, L. Kreplak, S.V. Strelkov, U. Aebi, Intermediate filaments: from cell architecture to nanomechanics, Nat Rev Mol Cell Biol 8(7) (2007) 562–73.

