## Supplementary Information for "Mimicking two posttranslational modifications associated with oxidative stress affords phase separation of vimentin"

Supplementary table 1
Supplementary figure 1

**Suppl. Table 1. Oligonucleotides for site directed mutagenesis**

| **Mutation** | **Template** | **Sequence 5' - 3'** |
|---|---|---|
| S49A | vim | GCCCCAGCACCGCCCGCAGCCTCTAC |
| S49D | vim | CGCCCCAGCACCGACCGCAGCCTCTAC |
| S56A | vim | CTCTACGCCTCGGCACCGGGCGGCGTGTATG |
| S56D | vim | CTCTACGCCTCGGACCCGGGCGGCGTGTATG |
| S72A | vim | CCGTGCGCCTGCGGGCCAGCGTGCCCGGGG |
| S72D | vim S72A | GTGCGCCTGCGGGACAGCGTGCCCGGGGTG |

Nucleotides mutated with respect to the template are shown in red

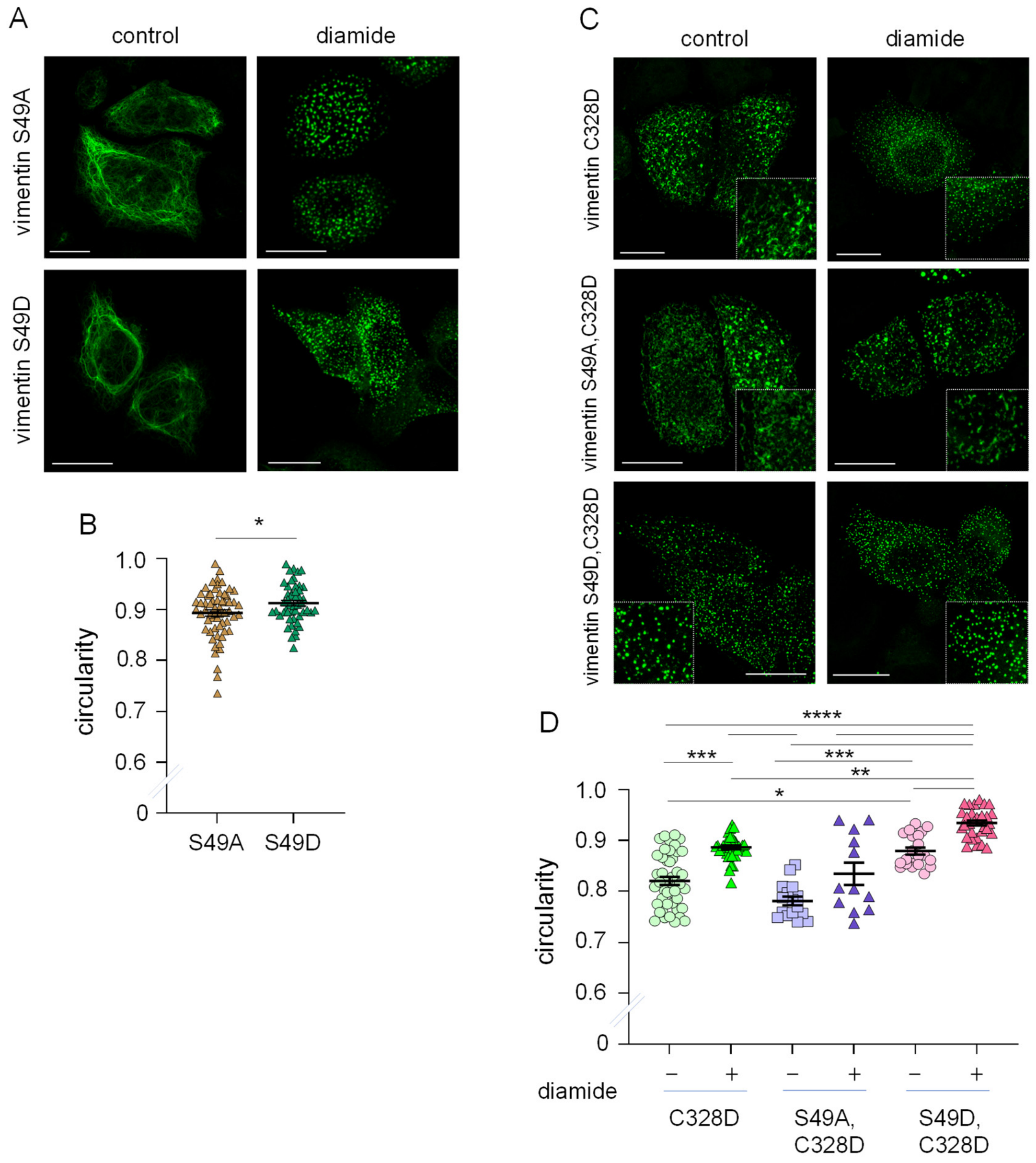


**Suppl. Fig. 1. Impact of mutations at the 49 site of vimentin on cellular assembly.** (A) SW13/cl.2 cells were transiently transfected with plasmids RFP//vimentin S49A or S49D and the distribution of the protein constructs upon 15 min treatment in the absence or presence of 1mM diamide was evaluated by immunofluorescence. (B) The circularity of the droplets formed by diamide treatment of cells expressing vimentin S49A or S49D was analyzed in at least 49 cells per experimental condition and the average ± SEM is represented. *p<0.05 by Student's t-test. (C) Cells were transiently transfected with plasmids RFP//vimentin C328D, RFP//vimentin S49A,C328D or RFP//vimentin S49D,C328D, and the distribution of protein constructs under control conditions or after diamide treatment was assessed as above. Insets show enlarged areas of interest. (D) The circularity of the assemblies formed in (C) was measured. Graph shows average values ± SEM of a total of at least 12 cells per condition, from at least 3 experiments. Statistical significance was evaluated by the Kruskal-Wallis followed by the Dunn's test for multiple comparisons. *p<0.05; **p<0.01; ***p<0.001; ****p<0.0001.
